# Atmospheric hydrogen oxidation extends to the domain archaea

**DOI:** 10.1101/2022.12.13.520232

**Authors:** Pok Man Leung, Rhys Grinter, Eve Tudor-Matthew, Luis Jimenez, Han Lee, Michael Milton, Iresha Hanchapola, Erwin Tanuwidjaya, Hanna A. Peach, Carlo R. Carere, Matthew B. Stott, Ralf B. Schittenhelm, Chris Greening

## Abstract

Diverse aerobic bacteria use atmospheric hydrogen (H_2_) and carbon monoxide (CO) as energy sources to support growth and survival. Though recently discovered, trace gas oxidation is now recognised as a globally significant process that serves as the main sink in the biogeochemical H_2_ cycle and sustains microbial biodiversity in oligotrophic ecosystems. While trace gas oxidation has been reported in nine phyla of bacteria, it was not known whether archaea also use atmospheric H_2_. Here we show that a thermoacidophilic archaeon, *Acidianus brierleyi* (Thermoproteota), constitutively consumes H_2_ and CO to sub-atmospheric levels. Oxidation occurred during both growth and survival across a wide range of temperatures (10 to 70°C). Genomic analysis demonstrated that *A. brierleyi* encodes a canonical carbon monoxide dehydrogenase and, unexpectedly, four distinct [NiFe]-hydrogenases from subgroups not known to mediate aerobic H_2_ uptake. Quantitative proteomic analyses showed that *A. brierleyi* differentially produced these enzymes in response to electron donor and acceptor availability. A previously unidentified group 1 [NiFe]-hydrogenase, with a unique genetic arrangement, is constitutively expressed and upregulated during stationary phase and aerobic hydrogenotrophic growth. Another archaeon, *Metallosphaera sedula*, was also found to oxidize atmospheric H_2_. These results suggest that trace gas oxidation is a common trait of aerobic archaea, which likely plays a role in their survival and niche expansion, including during dispersal through temperate environments. These findings also demonstrate that atmospheric H_2_ consumption is a cross-domain phenomenon, suggesting an ancient origin of this trait, and identify previously unknown microbial and enzymatic sinks of atmospheric H_2_ and CO.

## Introduction

Over the last 15 years it has been established that aerobic bacteria residing in soils and other aerated ecosystems are the primary biogeochemical sink of the global hydrogen (H_2_) cycle (1). These bacteria consume atmospheric hydrogen gas (tropospheric concentration ~0.53 ppmv / ~0.4 nM) (2) using high-affinity [NiFe]-hydrogenases from the subgroups 1h, 1l, 1f and 2a (3–7). They relay H_2_-derived electrons primarily to the aerobic respiratory chain, enabling ATP synthesis, though some bacteria also use them to sustain CO_2_ fixation (1, 5, 8). Diverse organoheterotrophic bacteria depend on atmospheric H_2_ as an energy source to survive during organic carbon limitation, as confirmed by genetic studies showing high-affinity hydrogenases enhance the long-term survival of *Streptomyces avermitilis* exospores (3, 9) and *Mycobacterium smegmatis* cells (10–12). Some bacteria also mixotrophically grow by co-oxidizing atmospheric H_2_ with other energy sources, as inferred from studies on the organotroph *Sphingopyxis alaskensis* (13), methanotroph *Methylocapsa gorgona* (14), nitrite oxidizer *Nitrospira moscoviensis* (15), and iron oxidizer *Acidithiobacillus ferrooxidans* (16). In temperate soils, genome-resolved metagenomic analysis suggests 32% of bacterial cells and at least 21 phyla encode hydrogenases to oxidize atmospheric H_2_ (17, 18). Moreover, atmospheric H_2_ oxidation sustains the energy, carbon, and hydration needs of bacteria in ecosystems with low primary production, such as Antarctic desert soils (7, 19). Atmospheric carbon monoxide (CO; tropospheric concentration ~0.09 ppmv / 0.086 nM) (20) is also a vital energy source sustaining the survival of aerobic bacteria (21, 22). Similar to atmospheric H_2_, this gas is oxidized by high-affinity form I CO dehydrogenases and the derived electrons are transferred to terminal oxidases (21–23). Overall, atmospheric H_2_ and CO both constitute dependable lifelines for bacterial survival given their ubiquity, diffusivity, and high-energy content (24).

To date, atmospheric H_2_ oxidation has not been observed in the domain archaea. Aerobic archaea are abundant community members in oxic environments, accounting for 1 – 5% in surface soils (25, 26) and 2 – 20% in ocean waters (27). In extreme environments such as oxic subseafloor sediments, acid mine drainage, salt lakes, and hot springs, aerobic archaea are relatively enriched and can constitute over half the microbial population (28, 29). Following the seminal report of aerobic H_2_ oxidation by thermoacidophilic archaea 30 years ago by Stetter *et al*. (30), numerous other isolates from geothermal habitats associated exclusively with Thermoproteota order Sulfolobales have been reported to grow aerobically on H_2_ as the sole electron donor but at elevated (above micromolar) concentrations (31–34). The enzymes responsible for this process were not resolved. A study on the thermoacidophile *Metallosphaera sedula* (Sulfolobales) demonstrated a [NiFe]-hydrogenase containing two hypothetical genes (*isp1*, *isp2*) was upregulated during aerobic autotrophic growth (35), though other studies have suggested this enzyme is oxygen-sensitive, and instead, involved in anaerobic sulfidogenic growth (36, 37). In light of recent discoveries in bacteria, it also remains to be tested whether archaea are capable of consuming H_2_ gas at sub-micromolar concentrations. Aerobic CO oxidation has been observed in thermoacidophilic (Sulfolobales) (38, 39) and halophilic (Halobacteria) (40, 41) archaea, the latter to sub-atmospheric levels (40, 41). While genomic and biochemical evidence suggests that this process is mediated by form I CO dehydrogenases, the physiological role of this process remains unresolved. Since atmospheric substrates are readily available to archaea living in oxic environments, a reasonable hypothesis is that some aerobic archaea use atmospheric H_2_ and CO to conserve energy during growth and survival akin to bacteria.

Here we studied the aerobic archaeon *Acidianus brierleyi* (DSM 1651^T^), which was isolated from an acidic hot spring in 1973 (42). This organism grows lithoautotrophically on micromolar levels of H_2_ using either elemental sulfur under anoxic conditions or oxygen under oxic conditions as terminal electron acceptors (30, 43). Its capacity to use CO as a substrate has not been tested. Consistent with the hyperthermoacidophilic growth of other members within Sulfolobales, *A. brierleyi* grows between 45 to 75°C (T_opt_ 70°C) at pH values of 1 to 6 (pH_opt_ 1.5 to 2.0) (43). Previous studies have demonstrated that geothermal isolates from the phyla Acidobacteriota (6, 44), Chloroflexota (45), Firmicutes (46), and Verrucomicrobiota (47) can each meet their energy needs using atmospheric H_2_. We therefore hypothesised that some H_2_-oxidizing archaea from such habitats, as represented by *A. brierleyi*, also use this gas. In this study, we characterized the kinetics, threshold, and temperature dependence of aerobic H_2_ uptake in *A. brierleyi* using ultra-sensitive gas chromatography. We also tested whether another hydrogenotrophic archaeon, *Metallosphaera sedula* (DSM 5348^T^), consumes atmospheric H_2_. To identify the enzymatic mediators of this process, we analysed the phylogeny, genetic organisation, and predicted protein complex structures of the hydrogenases of *A. brierleyi*. Finally, we performed quantitative proteomics analysis of *A. brierleyi* cells grown at various growth stages with different electron acceptors to substantiate these inferences and gain insights into the ecophysiological roles of H_2_ oxidation.

## Results

### *Acidianus brierleyi* rapidly oxidizes atmospheric H_2_ and CO across a range of temperatures

To determine if *A. brierleyi* is a high-affinity H_2_ oxidizer, we grew triplicate cultures organoheterotrophically in closed serum vials to mid-exponential growth phase (OD_600_ ~ 0.06; **Fig. 1A**), amended the vials with a dilute initial concentration of H_2_ (~ 10 ppmv / ~ 6 nM), and monitored H_2_ oxidation *via* gas chromatography. The cultures aerobically oxidized H_2_ in an apparent first-order kinetic process, reaching a sub-atmospheric threshold concentration of 0.20 to 0.25 ppmv (0.12 to 0.16 nM) within 24 hours (**Fig. 1B**). Concomitantly, there was no decrease of H_2_ in heat-killed cells and sterile medium, validating the observation is due to archaeal activity (**Fig. 1B, Table S1**). To extend these findings to other aerobic archaea, we performed the same experiment on another hydrogenotrophic Sulfolobales species, *M. sedula*. This archaeon consumed headspace H_2_ at a slower rate than *A. brierleyi*, but reached a lesser sub-atmospheric consumption threshold (~0.1 ppmv / 0.06 nM) during exponential growth (OD_600_ ~0.047) at 65°C (**Fig. 1B**). Together, these results provide the first evidence that atmospheric H_2_ oxidation is not exclusive to bacteria, but rather a cross-domain trait.

**Fig. 1.**
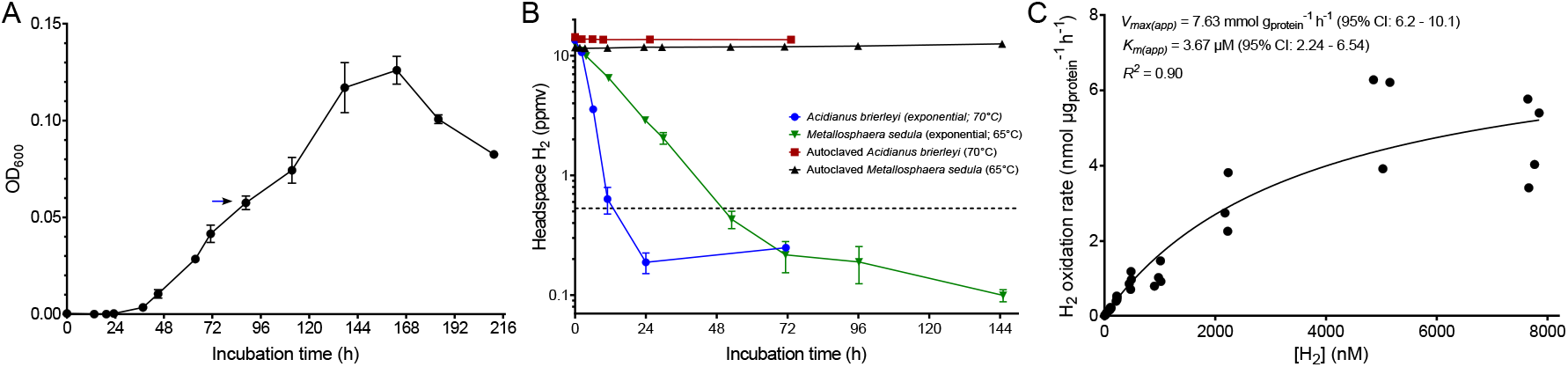
Aerobic hydrogen (H_2_) oxidation during exponential growth of *Acidianus brierleyi* at 70°C. **(A)** Heterotrophic growth of *A. brierleyi* in DSMZ medium 150 base with 0.2 g l^-1^ yeast extract at 70°C and pH 2. The blue arrow indicates the initial cell density where the H_2_ consumption experiment in (**B**) was performed on *A. brierleyi*. (**B**) Gas chromatography measurement of H_2_ oxidation to sub-atmospheric levels by *A. brierleyi* and *M. sedula* at mid-exponential growth, with heat-killed cells as negative controls. Headspace H_2_ mixing ratio is presented on a logarithmic scale and the dotted line indicates the mean atmospheric H_2_ mixing ratio (0.53 ppmv). Error bars in both (**A**) and (**B**) show one standard deviation of three biological replicates. (**C**) Apparent kinetics of H_2_ oxidation by mid-exponentially growing *A. brierleyi* cultures. A Michaelis–Menten non-linear regression model was used to calculate kinetic parameters and derive a best fit curve.

Next, we measured H_2_ uptake kinetics of in *A. brierleyi* whole cells during aerobic conditions. H_2_ uptake followed Michaelis-Menten kinetics (*R^2^* = 0.90) with an apparent half-saturation constant (*K*_m(app),70°C_) of 3.67 μM (2.24 - 6.54, 95% confidence interval) and an apparent maximum oxidation rate (*V*_max(app),70°C_) of 7.63 mmol g_protein_^-1^ h^-1^ (6.2 - 10.1, 95% confidence interval), respectively (**Fig. 1C**). It should be noted that these parameters potentially reflect the combined activity of two or more kinetically distinct hydrogenases within *A. brierleyi* cells, given hydrogenases with micromolar affinity constants typically cannot oxidize atmospheric H_2_ (5).

We then incubated mid-exponential and stationary phase cultures of *A. brierleyi* at 70°C with ~10 ppmv each of H_2_, CO, and methane (CH_4_; as an internal standard) in the ambient air headspace. Throughout the timecourse, there was minimal gas leakage reflected by CH4 concentrations within samples and controls (**Fig. S1**). A simultaneous decrease of headspace H_2_ and CO was observed in cultures under both conditions (**Fig. 2A**), confirming that the CO dehydrogenase is active and suggesting aerobic oxidation of both gases is a constitutive process. Whereas H_2_ was rapidly consumed to sub-atmospheric levels, CO was more slowly consumed to a supra-atmospheric steady-state concentration (1.3 – 2.1 ppmv). However, negative controls of sterile medium and killed cultures showed a linear and substantial production of CO (~ 2.0 ppmv day^-1^, *R^2^* = 0.99) (**Fig. 2A**), likely due to thermal degradation of the butyl-rubber stopper or organic substrates in the medium as previously observed (48,49).

**Fig. 2.**
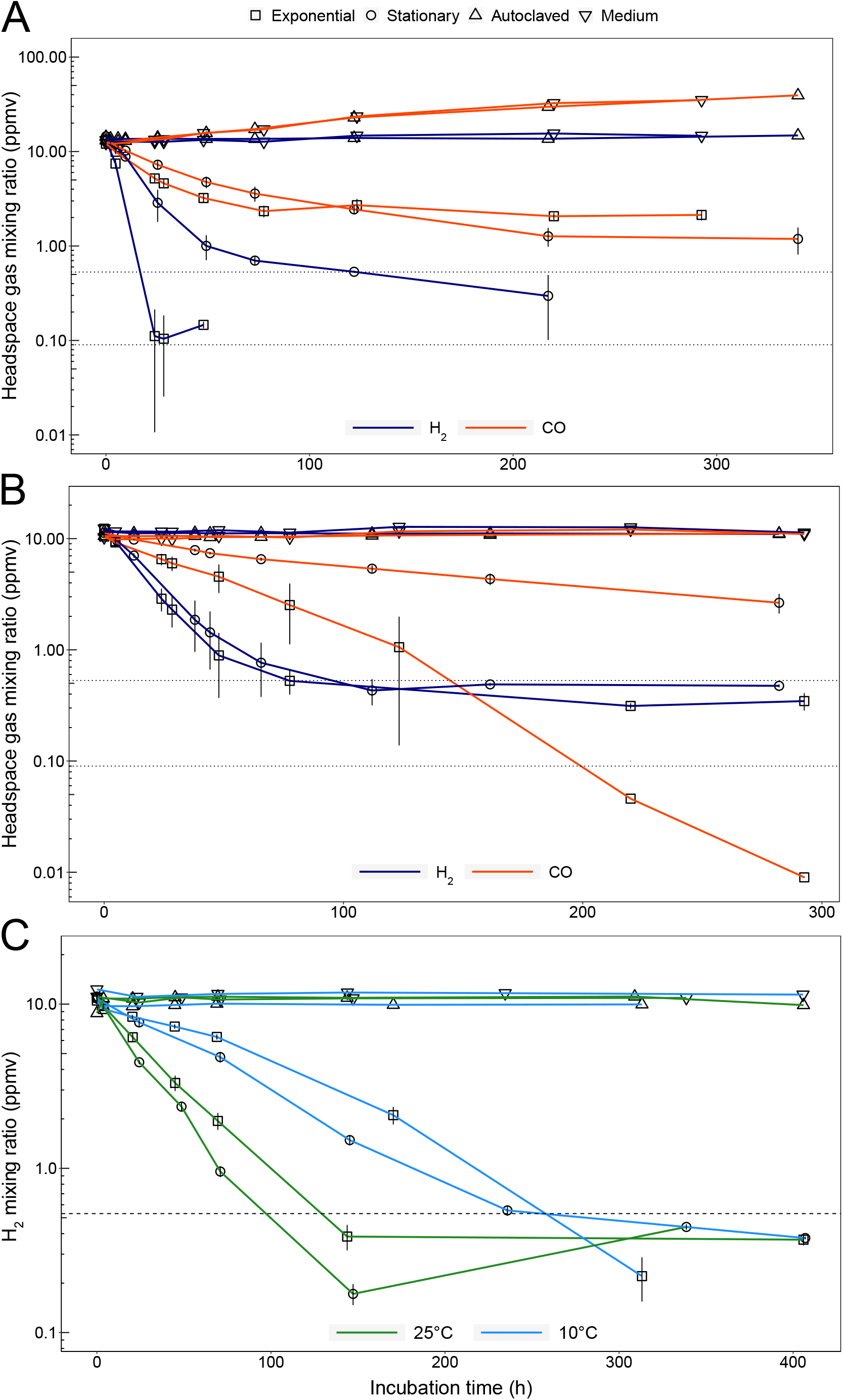
Co-oxidation of hydrogen (H_2_) and carbon monoxide (CO) by mid-exponential and stationary phase cultures of *Acidianus brierleyi* at various temperatures. Mid-exponential and stationary growth phase cells grown at 70°C were used for experiments (**Table S1**). Gas chromatography measurement of simultaneous consumption of H_2_ and CO oxidation at (**A**) 70°C and (**B**) 37°C. (**C**) Continued H_2_ oxidation activity at 25°C and 10°C. For all panels, error bars show one standard deviation of three biological replicates and the dotted lines indicate the mean atmospheric H_2_ (0.53 ppmv) and CO (0.09 ppmv) mixing ratios.

In order to mitigate this effect and establish an uptake threshold for CO, we incubated cultures at 37°C where abiotic CO production was 20 times slower (~0.10 ppmv day^-1^, *R^2^* = 0.50) (**Fig. 2B**). The cells co-consumed H_2_ and CO during the timecourse to below atmospheric levels (**Fig. 2B**). This result extends Thermoproteota as the second archaeal phylum capable of atmospheric CO oxidation in addition to Halobacteriota (40, 41). To determine if the species can oxidize trace gases at temperatures representative of temperate environments, H_2_ oxidation experiments were carried out at 25°C and 10°C. As with the 37°C experiments, cultures were first grown at 70°C to the desired density, then equilibrated and incubated at colder temperatures to measure gas consumption. Atmospheric H_2_ uptake was observed even two weeks after harvesting cultures (**Fig. 2C**), suggesting this process is highly resilient to temperature variations. Thus, an obligate thermophile *A. brierleyi* (T_opt_ 70 °C) continually harvests trace gases at temperatures outside its growth ranges (T_min_ 45°C) (42, 43).

### *Acidianus brierleyi* possesses four phylogenetically and syntenically distinct [NiFe]-hydrogenases widely distributed in Sulfolobales

We built an HMM profile using all hydrogenase large subunit reference sequences from HydDB (50) to comprehensively search for the enzymes responsible for H_2_ oxidation in *A. brierleyi*. Through careful inspection of hits, we identified four [NiFe]-hydrogenases encoded by this organism. They belong to uptake group 1 and 2 [NiFe]-hydrogenases but share < 30% sequence similarity with each other. To contextualize their relationships, a phylogenetic tree was constructed of the catalytic subunits of these hydrogenases, HydDB references, and newly identified group 1 and 2 [NiFe]-hydrogenases from all representative archaeal species in the Genome Taxonomy Database (GTDB) release 202 (51) (**Fig. 3 & Fig. S2**). The four hydrogenases clustered into distinct lineages. DFR85_RS19635 (HcaL) is a member of the previously defined 1g subgroup (18), also known as Crenarchaeaota Isp-type hydrogenase (52), characterized by the presence of genes encoding a putative transmembrane *b*-type cytochrome (*isp1*) and iron-sulfur protein (*isp2*) between the small and large subunit genes. This unique genetic arrangement is conserved in *A. brierleyi* (**Fig. 4A**). These enzymes are the sole biochemically and genetically characterized uptake [NiFe]-hydrogenase from Sulfolobales and are known to interact with sulfur reductase to mediate sulfidogenic H_2_ oxidation (36, 37, 53–55). They are also the only formally reported hydrogenase from *Acidianus*, and thus often assumed to mediate aerobic H_2_ oxidation (35). DFR85_RS29945 (HysL) clusters with subgroup 2e, a putative hydrogenase lineage defined through genome surveys (50) and only previously reported in *Metallosphaera sedula* (35). The gene cluster predicted to encode the structural subunits of this enzyme (*hysLSM*) is similar to the high-affinity group 2a enzyme from bacteria (**Fig. 4A**) (56).

**Fig. 3.**
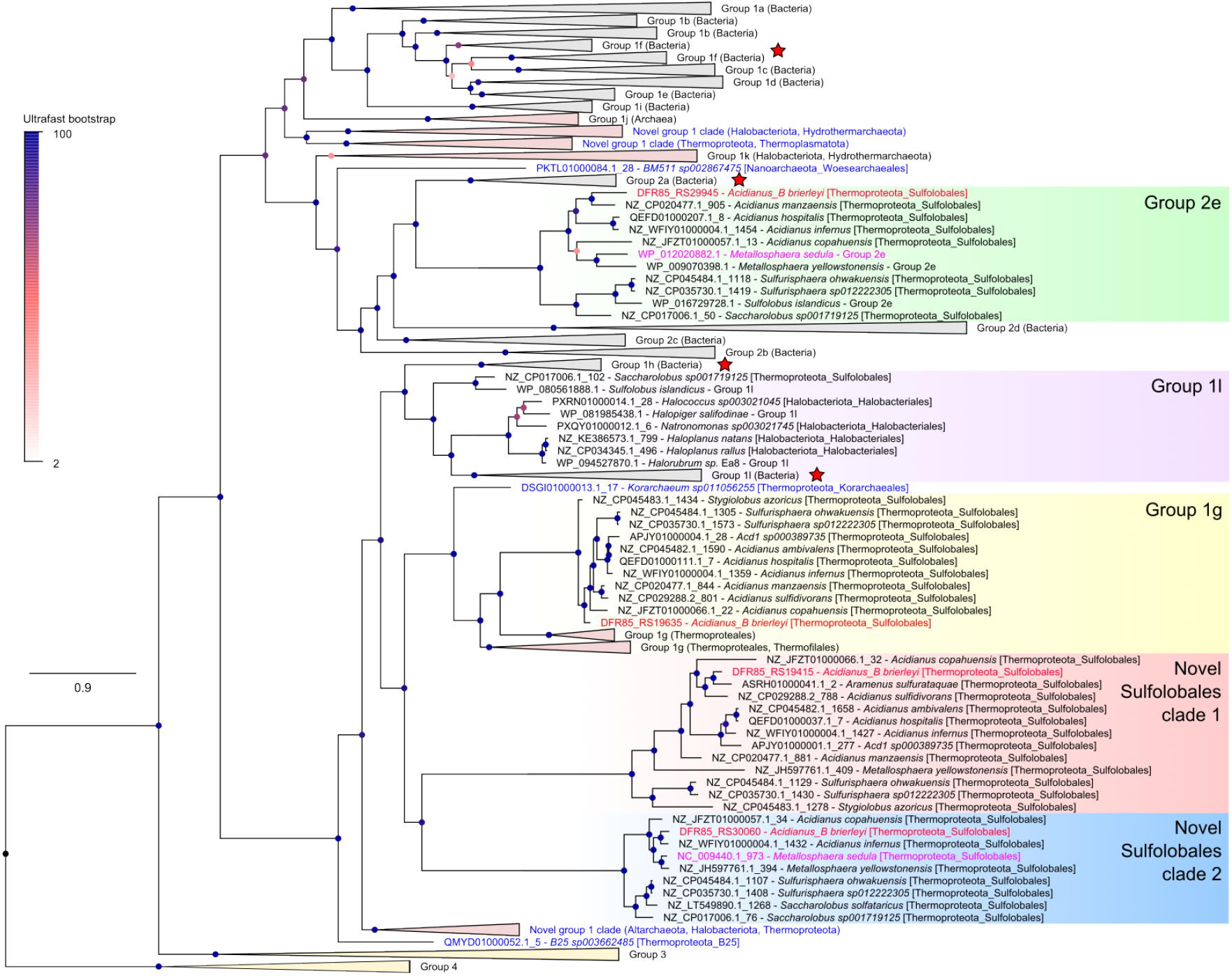
Identification of four [NiFe]-hydrogenases in *Acidianus brierleyi*. Maximum-likelihood phylogenetic reconstructions of amino acid sequences of the uptake group 1 and 2 [NiFe]-hydrogenase large (catalytic) subunits identified in *A. brierleyi* (four sequences; red text), *M. sedula* (two sequences; fuschia text), genomes of all archaeal representative species in Genome Taxonomy Database (GTDB) release 202 (202 sequences) (51), and hydrogenase reference database HydDB (1003 sequences) (50). Group 3 and 4 [NiFe]-hydrogenases were included as outgroups and the phylogeny was rooted between group 4 [NiFe]-hydrogenases and all other groups. Note that *A. brierleyi* is classified under a distinct genus from *Acidianus* (placeholder name *Acidianus_B*) in GTDB. Collapsed subgroups/clades that were exclusively bacterial or archaeal are shaded in grey or pink, respectively. Novel archaeal hydrogenase lineages are colored in blue or specified otherwise. Star symbols denote hydrogenase subgroups with members experimentally shown to mediate atmospheric H_2_ oxidation. Details on alignment and tree inference can be found in **Materials and Methods** and all sequences are provided in **Table S2**. Each node was colored by ultrafast bootstrap support percentage (1000 replicates) and the scale bar indicates the average number of substitutions per site. The tree showing all taxa is provided in **Fig. S2**.

**Fig 4.**
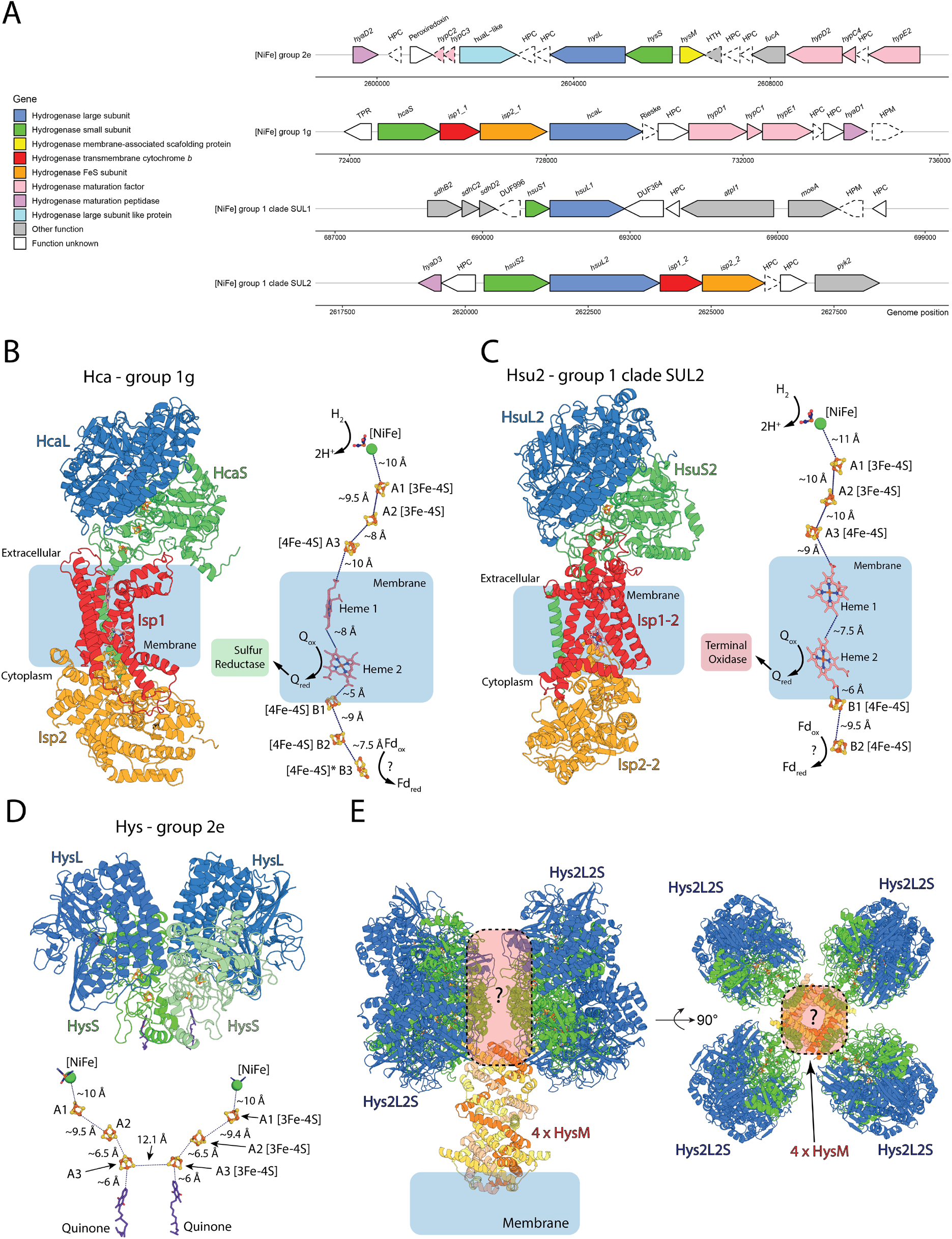
Genetic organization and AlphaFold structural models of the four [NiFe]-hydrogenases in *Acidianus brierleyi*. (**A**) Genetic organization of the four [NiFe]-hydrogenases encoded by *A. brierleyi*. Arrow outlines denote the presence (solid line) and absence (dotted line) of protein expression of the gene detected by shotgun proteomics under tested conditions. Gene length is shown to scale. HPC, hypothetical cytosolic protein; HPM, hypothetical membrane protein. Detailed information on loci, annotations and amino acid sequences of each gene are available in **Table S3**. AlphaFold-derived models of the group 1g hydrogenase Hca (**B**), the novel group 1 clade SUL2 hydrogenase Hsu2 (**C**), the group 2e hydrogenase Hys (**D**) shown as a cartoon representation of the complex formed by the subunits identified in panel A (**left/top panel**) and the electron transport relay formed by modelled cofactors, with predicted electron donors and acceptors indicated (**right/bottom panel**). (**E**) A model of a higher order complex formed by Hys, incorporating the AlphaFold model of the HysM subunit, and based on the Cryo-EM structure of the group 2a hydrogenase Huc from *Mycobacterium smegmatis* (60).

The two other deep-branching hydrogenases are from previously unidentified subgroups exclusive to Sulfolobales. Designated novel Sulfolobales clade SUL1 (HsuL1) and SUL2 (HsuL2) hereafter, they have the highest sequence similarity with the group 1l (31.4%; WP_080561888.1 - *Sulfolobus islandicus*) and 1h hydrogenases (29.3%; WP_015922819.1 - *Thermomicrobium roseum*) in HydDB respectively. SUL1 is encoded by a lone pair of large and small subunits. In contrast, the small and large subunit genes of SUL2 are immediately followed by *isp1* and *isp2* genes (**Fig. 4A**). This observation is unexpected given the *isp* genes were thought to be exclusive to two distantly related hydrogenase subgroups (bacterial 1e and archaeal 1g) and always encoded between the hydrogenase structural subunits (18, 37, 52, 57). SUL2 may thus represent a key ‘missing link’ to study the evolution of uptake hydrogenases. None of the four [NiFe]-hydrogenases affiliate directly with subgroups known to mediate atmospheric H_2_ oxidation (i.e. groups 1h, 1l, 1f, 2a), suggesting novel lineages catalyse this activity in *A. brierleyi*. Other novel uncharacterised hydrogenases were also widespread in other uncultured aerobic and anaerobic archaea. At least three other divergent and previously undescribed group 1 [NiFe]-hydrogenase clades, spanning Altarchaeota, Halobacteriota, Hydrothermarchaeota, Thermoproteota, and Thermoplasmatota, were also identified (**Fig. 3 & Fig. S2**).

To gain insight into the structure, subunit composition, and physiological functions of the four *A. brierleyi* hydrogenases, we modelled their structures using AlphaFold Multimer (58, 59). The results of this modelling are discussed in detail in **Supplemental Note 1** and summarised here. The models of both Hca (group 1g) and Hsu2 (clade SUL2) consist of extracellular large and small [NiFe]-hydrogenase subunits, which form a membrane-spanning complex with the integral membrane *b*-type cytochrome protein Isp1 associated with the multi iron-sulfur protein Isp2 on the cytoplasmic side of the membrane (**Fig. 4B,C**). This subunit arrangement gives rise to an electron transport relay that may allow both hydrogenases to transfer electrons from H_2_ oxidation to membrane-bound quinone via a heme group of Isp1 and to a lower potential electron acceptor (e.g. NAD^+^ or ferredoxin) in the cytoplasm via Isp2, possibly via electron bifurcation (**Fig. 4B,C**). This arrangement would thus enable these hydrogenases to provide electrons both to maintain the proton-motive force via quinone oxidation by either sulfur reductase or terminal oxidases, as well as drive carbon fixation during autotrophic growth. Modelling of Hys (group 2e) reveals an unexpected similarity to the recently characterized group 2a hydrogenase Huc from *Mycobacterium smegmatis*, which oxidizes H_2_ at sub-atmospheric concentrations and forms a large oligomer around the central membrane-associated subunit HucM (60). Despite limited sequence identity (~45%), modelling reveals the HysSL subunit forms a dimer, which is highly similar to that of HucSL (RMSD = 1.55 Å; **Fig. 4D**). In a similar manner, the modelled structure of HysM is a homologue to HucM (despite only sharing 18% sequence identity) and forms a characteristic tube-like structure, which in Huc scaffolds the enzyme and delivers quinone to the electron acceptor site of HucS (60). Based on these similarities, we generated a full model for Hys (**Fig. 4E**). Finally, the model for Hsu1 (clade SUL1) reveals a truncated HsuS1 subunit with a single [FeS]-cluster (**Fig. S3A**). Hsu1 may form a complex with an integral membrane quinone-reducing complex related to succinate dehydrogenase, encoded by genes directly upstream of HsuS1 (**Fig. S3B**).

Through analysis of the *A. brierleyi* genome, we also identified an operon encoding a form I CO dehydrogenase responsible for the observed carbon monoxide oxidation (**Fig. S4**). It has a typical genetic arrangement of form I CO dehydrogenase (*coxEDLSMF*), and its large subunit (*coxL*) harbours the conserved AYXCSFR active site signature motif (21). Phylogenetic analysis of the CoxL protein shows that the *A. brierleyi* enzyme clusters with those from diverse bacteria and other thermoacidophilic archaea, whereas halophilic archaea possess a distinct deeper-branching enzyme (22). Thus, members of the Sulfolobales likely acquired CO dehydrogenase from aerobic bacteria through horizontal gene transfer and independently of halophilic archaea.

### Quantitative proteome comparison identifies differentially regulated hydrogenases in response to principal growth substrates and terminal electron acceptors

To gain a system-wise understanding of how various hydrogenases contribute to *A. brierleyi* physiology and energetics, quadruplicate shotgun proteomes were compared for cells grown at four different conditions: mid-exponential growth on heterotrophic medium (EX); stationary phase on heterotrophic medium (ST); sulfur-dependent anaerobic hydrogenotrophic growth (H_2_:CO_2_:N_2_ = 20%:5%:75% v/v) on mineral medium (AN); and aerobic hydrogenotrophic growth (H_2_:CO_2_:air = 20%:5%:75% v/v) on mineral medium (AE). We have identified a total of 1847 proteins (~65% of protein-coding genes) falling below a predefined false discovery rate threshold of 1%. Principal component analysis showed that replicates were highly similar, but each sample group was distinct from one another (**Fig. S5**).

Proteome composition reflected the availability of the principal energy source and cellular status under each condition (**Fig. 5, Table S3**). Proteins involved in amino acid degradation, Entner-Doudoroff glycolytic pathway and tricarboxylic acid cycle for heterotrophic metabolism were highly abundant in heterotrophically grown cells (EX, ST) compared to cells at autotrophic conditions (AN, AE). For example, indolepyruvate:ferredoxin oxidoreductase subunit alpha (IorA) for amino acid degradation, 2-dehydro-3-deoxygluconokinase (KDGK) in the non-phosphorylative Entner-Doudoroff pathway, citrate synthase (GltA2), and succinate dehydrogenase flavoprotein subunit (SdhA) in the tricarboxylic acid cycle were significantly more abundant (6.7 – 21.6, 1.4 – 10.7, 3.2 – 5.5, and 1.8 – 2.2 fold respectively) under heterotrophic conditions (adj. *p* < 0.01) (**Fig. 5**). Contrastingly, cells under autotrophic conditions increased synthesis of proteins involved in 4-hydroxybutyrate cycle for carbon fixation and acetyl-CoA assimilation (adj. *p* < 0.001), including acetyl-CoA/propionyl-CoA carboxylase (AccADBC; 6 – 30 fold) and 4-hydroxybutyrate-CoA ligase (HbsC; 3.6 – 10.7 fold) (**Fig. 5, Table S3**). Furthermore, marker proteins involved in cell division (e.g. CdvA, Cdc48), translation (e.g. RLI, IF1A, EF2), amino acid biosynthesis (e.g. HisD, MetE, IlvC1), and nucleic acid biosynthesis (e.g. GuaA2, PyrG) were significantly reduced in stationary phase cultures in comparison to the other three actively growing cultures (adj. *p* < 0.01) (**Fig. 5**). Consistent with the activity-based measurements (**Fig. 2**), CO dehydrogenase subunits (CoxS, CoxM, CoxL) were produced at moderate levels during aerobic heterotrophic growth and high levels (average four-fold increase) during stationary phase (**Table. S3**). This is reminiscent of observations that CO dehydrogenase is upregulated and active during starvation in various bacterial cultures (22, 45, 61–63). Stationary phase cells also synthesized a greater abundance of extracellular solute-binding proteins (e.g. DppA1, DppA2, ESB2) and major facilitator superfamily sugar/acid transporters (MSAT) (**Fig. 6**, **Fig. S7**), likely enhancing scavenging of trace carbon substrates during energy limitation. Altogether, these results suggest that proteomics provides a reliable representation of cellular metabolism.

**Fig. 5.**
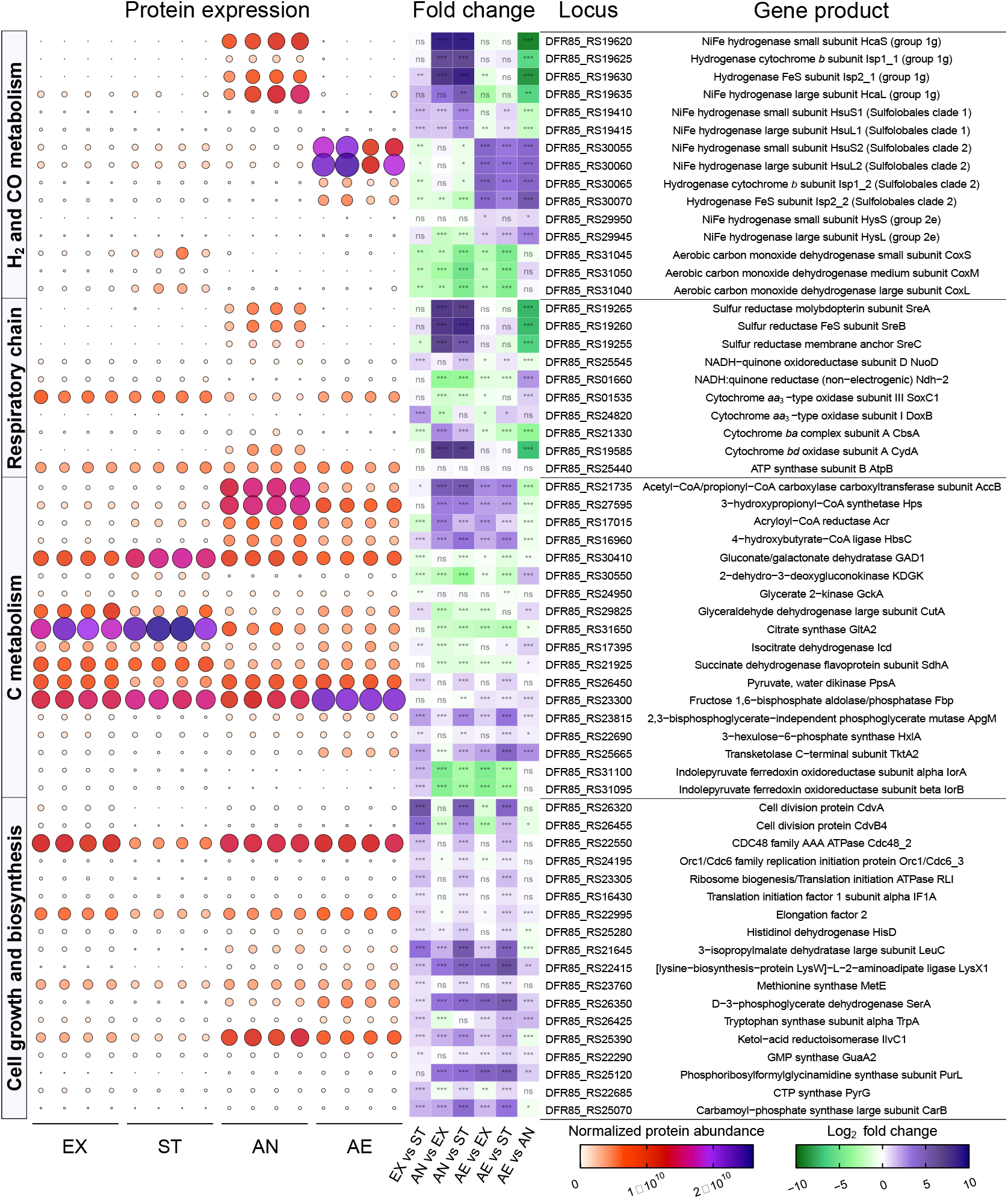
Quantitative comparison of selected *Acidianus brierleyi* proteins under heterotrophic growth, stationary phase, sulfur-dependent hydrogenotrophic growth and aerobic hydrogenotrophic growth. Culture condition (four biological replicates each): EX, mid-exponential growth phase on heterotrophic medium; ST, stationary phase on heterotrophic medium; AN, anaerobic sulfur-dependent hydrogenotrophic growth; AE, aerobic hydrogenotrophic growth (**Methods and Materials**). Normalised protein abundance value represents MaxLFQ total intensity for the protein. Bubble size and color indicate protein abundance of the corresponding gene product in each biological replicate. Significant difference in fold changes of protein abundance of each condition pair is denoted by asterisks (adjusted p value ⩽0.001, ***; ⩽0.01, **; ⩽0.05, *; > 0.05, ns). The full set of quantitative proteomics results is provided in **Table S3**.

**Fig. 6.**
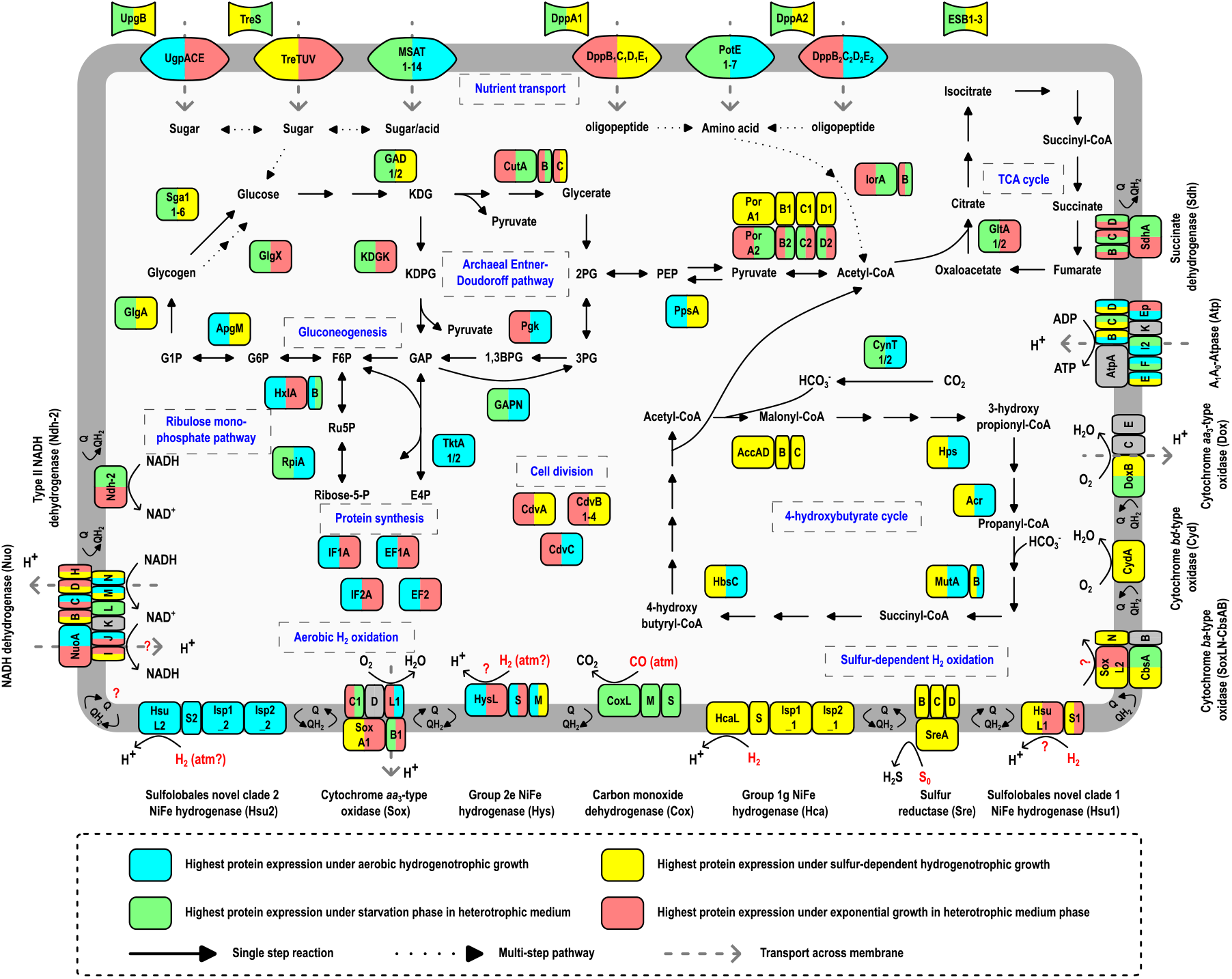
Genome and proteome-based model of H_2_ metabolism in heterotrophic and hydrogenotrophic growth of *Acidianus brierleyi*. The color scale indicates the conditions where proteins had the highest expression (left) and second highest expression if their log2 fold difference is less two (right). Metabolic marker genes for central carbon metabolism, trace gas oxidation, and respiratory chain are shown. AccADBC, acetyl-CoA/propionyl-CoA carboxylase; Acr, acryloyl-CoA reductase; ApgM, phosphoglycerate mutase; CdvABC, cell division proteins; CutABC, glyceraldehyde dehydrogenase; CynT, carbonic anhydrase; DppBCDE, ABC di/oligopeptide transporter; EF1A/EF2; elongation factors; GAD, gluconate/galactonate dehydratase; GAPN, glyceraldehyde-3-phosphate dehydrogenase; GlgA, glycogen synthase; GlgX, glycogen debranching protein; GltA, citrate synthase; HbsC, 4-hydroxybutyrate-CoA ligase; Hps, 3-hydroxypropionyl-CoA synthetase; HxlA, 3-hexulose-6-phosphate synthase; HxlB, 6-phospho-3-hexuloisomerase; IF1A/IF2A, translation initiation factors; IorAB, Indolepyruvate:ferredoxin oxidoreductase; KDGK, 2-dehydro-3-deoxygluconokinase; MSAT, major facilitator superfamily sugar/acid transporter; MutAB, methylmalonyl-CoA mutase; Pgk, phosphoglycerate kinase; PorABCD, pyruvate synthase; PotE, APC amino acid permease; PpsA, pyruvate, water dikinase; RpiA, ribose 5-phosphate isomerase; Sga1, glucoamylase; TktA, transketolase; UgpACE/TreTUV, ABC sugar transporter; UpgB/TreS/DppA/ESB, extracellular solute-binding proteins. Note that *A. brierleyi* is also known to grow chemolithoautotrophically on various reduced sulfur compounds and Fe^2+^ (**Fig. S7**), as extensively described in previous studies (55, 101, 102).

For proteins involved in H_2_ metabolism, the regulation of the group 1g [NiFe]-hydrogenase is in line with its reported role in sulfur-dependent hydrogenotrophy. Subunits of this hydrogenase (HcaS1, Isp2_1, Isp1_1) and sulfur reductase (SreA, SreB, and SreC) were among the ten most upregulated proteins during anaerobic autotrophic growth (adj. *p* < 0.001) (Fig. 5, Table S3). These proteins were otherwise undetected in the three oxic conditions (except lowly present in a single replicate of AE). HcaS1L1 represents 87.5% of all hydrogenase large and small subunits during anaerobic autotrophic growth, but has the lowest abundance during aerobic autotrophic growth. Hydrogenase maturation factors (*hypD1*, *hypC1*, *hypE1*) and proteases (*hyaD1*) neighbouring this hydrogenase shared a similar expression profile (Fig. S6). Expression of the novel hydrogenase lineage SUL1 is similar to Hca, as HsuL1 and HsuS1 had their highest abundance under anaerobic autotrophic growth, though only constituted 3.3% of hydrogenase proteins at this condition (Fig. 5). In contrast, the novel hydrogenase lineage SUL2 displayed a reverse expression pattern in which its abundance peaked during aerobic hydrogenotrophic growth (Fig. 5, Fig.6) compared to the three other conditions (adj. *p* < 0.001) (Fig. 5, Table S3). Intriguingly, SUL2 is the only hydrogenase that was found to be constitutively and copiously produced at all four tested conditions. HsuL2 and HsuS2 represents 58.4%, 79.2%, 9.1%, and 97.6% of all quantified hydrogenase large and small subunits during exponential heterotrophic, stationary heterotrophic, anaerobic autotrophic, and aerobic autotrophic conditions respectively (Table S3). Comparing expression in heterotrophically grown cultures, this hydrogenase was upregulated in the carbon-limited stationary phase condition in a manner reminiscent of most high-affinity group 1h [NiFe]-hydrogenases studied in bacteria (24). Finally, the group 2e hydrogenase had the lowest abundance among the four hydrogenases in all tested conditions (Fig. 5). Contrary to previous speculation, it is therefore unlikely to be involved in aerobic hydrogenotrophic growth (50), though may still be expressed at sufficient levels to scavenge atmospheric H_2_.

## Discussion

In this work, we report that two aerobic archaea scavenge H_2_ during growth and starvation, including at sub-atmospheric levels. These observations expand the guild of atmospheric H_2_ oxidizers to include domain archaea and suggest that this domain enhances the climate-relevant biogeochemical H_2_ sink. The aerobic oxidation of H_2_ by *A. brierleyi* follows Michaelis-Menten kinetics with a high K_m(app),70°C_ in micromolar ranges of H_2_ but a low uptake threshold of sub-nanomolar H_2_, highlighting that the organism efficiently uses a very broad range of H_2_ concentrations. The constitutive production and kinetic properties of the hydrogenases of *A. brierleyi* likely confer competitive advantages in dynamic geothermal habitats. Cells can rapidly mobilise H_2_ available at elevated concentrations, for example, from volcanic efflux and venting steam, or at oxic-anoxic interfaces where fermentative H_2_ is produced and diffuses from deeper anoxic layers (64, 65), to fuel autotrophic or mixotrophic growth. During periods when substrates are variable and limiting, the ability to harvest low concentrations of H_2_ either from the source environment or the atmosphere, for example, in the immediate vicinity of air-sediment or air-water interfaces, will enable them to meet maintenance energy requirements. This supplemental energy source can be particularly beneficial for organisms living in acidic and heated environments, as continuous energy expenditure is required to maintain a neutral intracellular pH against a steep external proton gradient and synthesise defence biomolecules to protect against oxidative and thermal damage (66, 67). Apart from demonstrating *A. brierleyi* is an atmospheric H_2_ oxidizer, we additionally confirm that this archaeon is an atmospheric CO oxidizer, extending this metabolism to a second archaeal phylum (Thermoproteota). Reflecting CO dehydrogenase is most highly expressed in stationary phase cultures, this metabolism may primarily support survival during organic carbon starvation, in line with observations in bacteria (22, 45, 61–63).

While aerobic oxidation of H_2_ in Sulfolobales has been widely reported, it is likely that previous studies misattributed the enzyme responsible for this metabolism. Group 1g [NiFe]-hydrogenases are the only hydrogenases previously characterized in this order, with group 2e enzymes also reported based on comparative genomic analysis (18, 35). However, these enzymes are known to couple H_2_ oxidation with sulfur reduction (but not with oxygen), and are commonly present in obligately anaerobic archaea. In this study, we revealed the presence of two additional putative uptake [NiFe]-hydrogenases in *A. brierleyi* and the wider Sulfolobales order (**Fig. 3**), designated SUL1 and SUL2. Together with comprehensive proteomic analysis, we introduce a model of H_2_ metabolism in heterotrophic and hydrogenotrophic growth of *A. brierleyi* (**Fig. 6**). Reflecting the high abundance during sulfur-dependent hydrogenotrophic growth but the minimal expression under oxic conditions, the group 1g [NiFe]-hydrogenase functions as the primary hydrogenase for H_2_ oxidation under anaerobic conditions and relays electrons to sulfur as the terminal electron acceptor. Given its overwhelming expression during aerobic hydrogenotrophic growth, the novel hydrogenase SUL2 is the predominant hydrogenase for H_2_ oxidation under aerobic conditions and transfers electrons to oxygen as the terminal electron acceptor. In agreement with this notion, homologs of SUL2 and group 2e hydrogenases are the sole hydrogenases identified in the aerobic hydrogenotroph *M. sedula* (30) (**Fig. S2**). Some electrons derived from both hydrogenases are potentially transferred to NADH dehydrogenase through reverse electron transport to generate reductant necessary for carbon fixation via the 4-hydroxybutyrate cycle. Alternatively, as inferred from AlphaFold structural modelling (**Fig. 4**), the SUL2 and group 1g enzymes may form electron-bifurcating complexes that relay electrons from H_2_ to high-potential quinone through Isp1 subunit for the proton motive force generation and low-potential acceptors (e.g. ferredoxin) through Isp2 subunit for carbon fixation. This efficient energy conservation mechanism may explain the dominant roles of both hydrogenases during hydrogenotrophic growth. Direct experimental validation using purified enzymes would be necessary to confirm this.

Although proteomic evidence strongly suggests hydrogenase SUL2 is responsible for aerobic oxidation of H_2_ at high concentrations, it is unresolved whether this enzyme mediates atmospheric uptake. Based on our result that the obligately aerobic *M. sedula* is a high-affinity H_2_ oxidizer (**Fig. 1B**), it is logical to constrain the enzyme mediating atmospheric H_2_ oxidation to the only two hydrogenases it encodes, namely the SUL2 or group 2e [NiFe]-hydrogenases (**Fig. 3**). SUL2 potentially mediates atmospheric H_2_ oxidation based on four lines of evidence: (i) it is the only hydrogenase constantly detected and abundantly expressed in all tested conditions, in agreement with the constitutive activity of atmospheric H_2_ oxidation in oxic conditions (**Fig. 2**); (ii) it shares the highest similarity with group 1h [NiFe]-hydrogenases known to mediate atmospheric H_2_ uptake in bacteria, albeit at < 30% identity; (iii) the lineage is only present in aerobic or facultatively aerobic Sulfolobales species (**Fig. 3**); and (iv) its high expression during stationary phase (compared to exponential phase) is analogous to the regulation of many group 1h [NiFe]-hydrogenases (**Fig. 5**) (24). The micromolar whole-cell H_2_ affinity constant is more consistent however with low-affinity [NiFe]-hydrogenases being the dominant H_2_-metabolising enzymes (5). The group 2e [NiFe]-hydrogenase may alternatively account for atmospheric H_2_ uptake. This reflects that this enzyme is a direct sister lineage and shares a highly similar predicted protein structure to the high-affinity bacterial group 2a hydrogenase (**Fig. 3, Fig. 4**), is solely present in aerobic or facultatively aerobic Sulfolobales species (**Fig. 3**), and is downregulated under anaerobic conditions (**Fig. 5**). Though it is expressed at relatively low levels under tested conditions, such levels may be sufficient for a catalytically efficient high-affinity hydrogenase to conserve sufficient energy from atmospheric H_2_. Indeed, the wide kinetic range of *A. brierleyi* could be explained through the complimentary activity of both low-affinity (e.g. SUL2) and high-affinity (e.g. group 2e) enzymes. The novel hydrogenase SUL1 is unlikely to account for atmospheric H_2_ oxidation given its presence in the obligately anaerobic *Stygiolobus azoricus* (68) (**Fig. 3**), decreased abundance in oxic conditions (**Fig. 5**), and absence in *M. sedula*. Heterologous expression in genetically tractable thermoacidophiles such as *Sulfolobus acidocaldarius* and *Sulfolobus islandicus* offers a possible means to resolve the function of each hydrogenase (69).

Our results also show that CO and H_2_ oxidation can co-occur at temperatures below growth ranges of extremely thermoacidophilic *A. brierleyi*. H_2_ oxidation remained active at temperatures as cold as 10°C for over two weeks, although future studies are needed to determine the lowest uptake temperature thresholds. In microbial ecology, the mechanisms of dispersal of extremophilic microorganisms such as nonsporulating obligate thermoacidophiles remain a long-standing mystery. Microorganisms that are taxonomically and genomically closely related are often found across geographically distant and disconnected thermal habitats (70, 71). Despite theory on a strong geographic barrier and endemism (72, 73), pioneer populations must be seeded from somewhere. How could these habitat specialists survive across temperate environments to distant specialised habitats? Here we propose hidden metabolic flexibility enables cells to meet energy needs during dispersal. During the dispersal and transition throughout temperate environments, trace gas oxidizers such as *A. brierleyi* may rely on the continual acquisition of atmospheric H_2_ and CO as universal metabolizable substrates for persistence. This may also help explain why terrestrial geothermal habitats select for diverse high-affinity trace gas oxidizers, including from the genera *Pyrinomonas* (6, 44), *Chloroflexus* (16), *Kyrpidia* (46), *Thermomicrobium*, *Thermogemmatispora* (16, 45), and *Methylacidiphilum* (47), despite being typically enriched with geothermally derived H_2_.

Lastly, atmospheric H_2_ oxidation may be a wider metabolic trait among aerobic archaea than previously assumed. The full set of four hydrogenases encoded by *A. brierleyi* are often possessed by other Sulfolobales members, including from the genera *Acidianus*, *Metallosphaera*, and *Sulfurisphaera* (**Fig. 3**). Certain halophilic (e.g. *Halopiger salifodinae*) and thermophilic (e.g. *S. islandicus* strain M.16.2) archaea encode [NiFe]-hydrogenases clustered with the high-affinity group 1h and 1l subgroups (**Fig. 3**). It is very probable that at least some aerobic archaea consume atmospheric H_2_ through these enzymes. Altogether, these findings suggest that atmospheric H_2_ oxidation is mediated by a broader range of microorganisms and enzymes than previously realised, and has a deeper-rooted evolutionary origin than initially hypothesised (74).

## Methods and Materials

### Archaeal strain and growth conditions

Pure cultures of the thermoacidophilic archaeal strains *Acidianus brierleyi* (DSM 1651^T^) and *Metallosphaera sedula* (DSM 5348^T^) were imported from Leibniz Institute DSMZ - German Collection of Microorganisms and Cell Cultures (DSMZ) in November 2021 and October 2022, respectively. *A. brierleyi* was regularly maintained in DSMZ medium 150, except that elemental sulfur was excluded. Per litre water, modified DSMZ medium 150 has a composition of 3 g (NH_4_)2SO_4_, 0.5 g K_2_HPO_4_ • 3 H_2_O, 0.5 g MgSO_4_ • 7 H_2_O, 0.1 g KCl, 0.01 g Ca(NO_3_)_2_, and 0.2 g yeast extract, with pH adjusted to 1.5 - 2.5 by 10N H_2_SO_4_ before autoclaving. *M. sedula* was regularly maintained heterotrophically in modified DSMZ medium 88. Per litre water, modified DSMZ medium 88 has a composition of 1.3 g (NH_4_)2SO_4_, 0.28 g KH_2_PO_4_, 0.25 g MgSO_4_ • 7 H_2_O, 0.07 g CaCl_2_ • 2 H_2_O, 0.02 g FeCl_3_ • 6 H_2_O, and 0.2 g yeast extract, with pH adjusted to 2 by 10N H_2_SO_4_ before autoclaving. Organic substrate stock solutions (10% w/v) were autoclaved separately before adding to the mineral medium. Liquid cultures of *A. brierleyi* (70 °C) and *M. sedula* (65 °C) were incubated with 100 rpm agitation in ambient air in the dark, unless otherwise specified. Cultures were propagated every 10 – 15 days with 1 - 10% v/v culture as inoculum in fresh medium pre-warmed at respective incubation temperatures.

### Growth characterization and biomass quantification

To characterize heterotrophic growth patterns of *A. brierleyi*, 1% v/v late exponential growing cultures were inoculated in 30 ml respective fresh medium in 125 ml aerated conical flasks. Triplicate cultures were then incubated at 70 °C with shaking at 100 rpm in the dark. Growth was monitored spectrophotometrically by measuring culture optical density at 600 nm (OD_600_; Eppendorf BioSpectrometer Basic) for up to six weeks. To quantify biomass of *A. brierleyi*, total cell protein was measured using the bicinchoninic acid protein assay (Sigma-Aldrich) against bovine serum albumin standards. Cell lysates for the assay were prepared as follows: 2 ml cultures were centrifuged at 20,200 × g for 30 min. The cell pellet was washed with 1 ml DSMZ medium 150 mineral base and centrifuged for 10 min, twice, to remove free proteins from the medium. The washed cell pellet was then resuspended in 500 μl of Milli-Q water and lysed through probe sonication (output setting: 4; 10 s twice; Branson Digital Sonifier 450 Cell Disruptor). Subsequently, the cell lysate was cooled down to room temperature and 100 μl of which was added to working reagents.

### Genome analysis

The closed genome and proteome annotations of *A. brierleyi* DSM 1651^T^ (75) were accessed through NCBI GenBank (76) under the assembly accession number GCA_003201835.2. Predicted protein coding sequences were further annotated against NCBI Conserved Domain Database (CDD) v3.19 (77) using rpsblast (-evalue 0.01 -max_hsps 1 - max_target_seqs 5) in BLAST+ v2.9.0 (78) and the Pfam protein family database v34.0 (79) using PfamScan v1.6 (default setting) (80). Protein subcellular localization and the presence of internal helices of the gene were predicted using PSORTb v3.0.3 (--archaea) (81). Metabolic pathway analysis was performed using DRAM v1.2.4 (82) with KEGG protein database (accessed 22 November 2021) (83). Catalytic subunits of [NiFe] hydrogenases were identified and classified using HydDB (50). The R package gggenes v0.4.1 (https://github.com/wilkox/gggenes) was used to construct gene arrangement diagrams. All sequences, annotations, and genetic arrangements are summarized in **Table S3**.

### Identification and phylogenetic analysis of group 1 and 2 [NiFe]-hydrogenase

All large subunit amino acid reference sequences of [NiFe]-hydrogenases were downloaded from HydDB (50), aligned using Clustal Omega v1.2.4 (84) and used to build a profile hidden Markov model (HMM) using HMMER v3.3.2 (85). The HMM profile was used to search against the open reading frames of *A. brierleyi* genome and all representative genomes from the Genome Taxonomy Database (GTDB) release 202 (51). Each match shorter than 100,000 residues and with a bit score ≥ 34.5 was retained and then filtered by the presence of both conserved proximal and medial CxxC motifs required to ligate [NiFe]-hydrogenase H_2_-binding metal centres (18, 86). The matches were also manually inspected through a combination of CDD annotations (77) and phylogenetic analysis. A multiple sequence alignment was performed on the retained archaeal and HydDB group 1 and 2 [NiFe]-hydrogenase sequences using MAFFT-L-INS-i v7.505 (87), with selected group 3 and 4 [NiFe]-hydrogenases as outgroups. The best-fit substitution model (Q.pfam+R10) was determined using ModelFinder (88) implemented in IQ-TREE2 v2.2.0 (89). A maximum likelihood phylogenetic tree was constructed using IQ-TREE2 v2.2.0 (89) with 1000 ultrafast bootstrap replicates (90).

### Computation modelling of hydrogenase complexes

[NiFe]-hydrogenase models were generated using AlphaFold multimer v2.1.1 implemented on the MASSIVE M3 computing cluster (58, 91). The amino acid sequences for the large and small subunits for each of the [NiFe]-hydrogenases identified in *A. brierleyi* (group 1g, group 2e, novel Sulfolobales clade 1 and 2) were modelled both alone and with the sequences of putative complex partners present in the hydrogenase gene cluster. Models produced were validated based on confidence scores (pLDDT), with only regions with a confidence score of > 85 utilized for analysis. Where complexes were predicted, subunit interfaces were inspected manually for surface complementarity and the absence of clashing atoms. Interfaces were also analyzed for stability using the program QT-PISA (92, 93), with only interfaces predicted to be stably utilized for analysis. To assign cofactors to the AlphaFold model subunits, the closest homologous structural domain for each modelled protein were identified by searching the PDB database using NCBI BLAST with the amino acid sequence of the protein as the query or the DALI server using the AlphaFold model as the query (94, 95). The homologous structures were aligned with the AlphaFold models in Pymol, and cofactors were added in corresponding positions to that of the experimental structures if all conserved coordinating residues were present. Cofactor position was then manually adjusted to optimize coordination and to minimize clashes. The following structures were used to model cofactors for the hydrogenases:

|  |  |  |  |  |  |
| --- | --- | --- | --- | --- | --- |
| *Group 1g* | *HcaS*<br>5AA5 chain A | *HcaL*<br>5AA5 chain C | *Isp1_1*<br>3EGW chain C | *Isp2_1*<br>7BKB chains P/Q |  |
| *Group 2e* | *HysS*<br>8DQV chain B | *HysL*<br>8DQV chain A | *HysM*<br>N/A |  |  |
| *SUL1* | *HsuS1*<br>4C3O chain B | *HsyL1*<br>4C3O chain B | *SdhB2*<br>1NEK chain B | *SdhC2*<br>1NEK chain C | *SdhD2*<br>1NEK chain D |
| *SUL2* | *HsuS2*<br>4C3O chain B | *HsuL2*<br>4C3O chain B | *Isp1_2*<br>3EGW chain C | *Isp2_2*<br>7BKB chains P/Q |  |

*PDB entries:*

- 5AA5 - Actinobacterial-type [NiFe]-hydrogenase from *Ralstonia eutropha*
- 3EGW - The crystal structure of the NarGHI mutant NarH
- 7BKB – Formate dehydrogenase-heterodisulfide reductase - formylmethanofuran dehydrogenase complex from *Methanospirillum hungatei*
- 8DQV - CryoEM structure of the [NiFe]-hydrogenase Huc from *Mycobacterium smegmatis*
- 4C3O - Structure and function of an oxygen tolerant [NiFe]-hydrogenase from *Salmonella*
- 3RGW - Crystal structure at 1.5 A resolution of an H_2_-reduced, O_2_-tolerant hydrogenase from *Ralstonia eutropha*
- 1NEK - Complex II (Succinate dehydrogenase) from *E. coli* with ubiquinone bound
- 3AYX - Membrane-bound respiratory [NiFe]-hydrogenase from *Hydrogenovibrio marinus*

### Hydrogen and carbon monoxide consumption measurement

Gas chromatography assays were carried out to test for the ability of *A. brierleyi* to oxidize H_2_ and CO at mid-exponential (OD_600_: 0.04 – 0.08 pre-OD_max_) and stationary growth phase (OD_600_: 0.04 – 0.08 post-OD_max_). 30 ml culture was transferred into a sterile 160-ml serum vial. Cultures were then allowed to adapt to the experimental conditions (70°C and agitation speed of 100 rpm) for at least 2 h before sealing the vial with butyl rubber stoppers. Butyl rubber stoppers throughout all trace gas experiments were pre-treated with 0.1 N hot NaOH solution, according to the methods of Nauer *et al*. (48), to decrease abiotic emissions of H_2_ and CO from the stopper. The ambient air headspace was amended with H_2_, CO and CH_4_ (via a mixed gas cylinder containing 0.1 % v/v H_2_, CO and CH_4_ each in N_2_, BOC Australia) to give initial mixing ratios of approximately 10 parts per million (ppmv) for each gas. CH_4_ was included as an internal standard to check for any gas leakage. To monitor changes in headspace gas concentrations, 2 ml of headspace gas was withdrawn using a gas-tight syringe at each sampling timepoint. A VICI gas chromatographic machine with a pulsed discharge helium ionization detector (model TGA-6791-W-4U-2, Valco Instruments Company Inc.) was then used to quantity H_2_ and CO concentrations. The machine was calibrated against ultra-pure H_2_ and CO standards down to the limit of quantification (H_2_: 20 ppbv; CO: 9 ppbv) (45). The machine was regularly calibrated against various calibration mixed gases with known trace gas concentrations. For *A. brierleyi*, the assays were also carried out at incubation temperatures of 10 °C, 25 °C (room temperature), and 37 °C to determine thermal plasticity of trace gas oxidation activities. For all conditions, a heat-killed culture (autoclaved at 121°C, 15 p.s.i. for 30 mins) and a medium only control were included. The same experimental set up was also prepared to measure H_2_ consumption by mid-exponentially growing *M. sedula* culture, except that the incubation temperature was at 65°C.

### *Acidianus brierleyi* hydrogen uptake kinetic analysis

Gas chromatography assays were used to determine uptake kinetics of H_2_ by *A. brierleyi*. Quadruplicate cultures at exponential growth phase were incubated independently with a wide range of starting headspace H_2_ concentrations (approximately 10, 20, 40, 80, 160, 320, 640, 1500, 3000, 6000, and 10000 ppmv) at 70 °C. Headspace H_2_ concentrations from 10 to 1500 ppmv and from 3000 to 10000 ppmv were amended via 1 % v/v H_2_ (in N_2_; Air Liquide) and ultra-high purity H_2_ (99.999%; BOC Australia), respectively. Quadruplicate medium only controls were also included for each concentration to monitor any gas leakage or sampling loss. Ten minutes prior to the first gas sampling, vial headspace overpressure was briefly released by a 22G needle within the incubator at 70 °C to equilibrate with atmospheric pressure. Headspace H_2_ concentrations were quantified at four time points, 0 h, 1 h, 2 h, and 4 h. To account for any potential lag response of the cultures to the concentrations of H_2_ supplemented and allow sufficient time for equilibration of the headspace H_2_ with the aqueous phase, reaction rates of H_2_ consumption were calculated based on the concentration change between the third and fourth time points. The dissolved concentrations of gases in aqueous phase at equilibrium state and at 1 atmospheric pressure were calculated according to Henry’s law and van’t Hoff equation as follows:

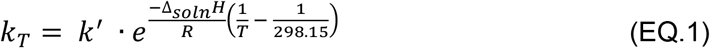

where *k_T_* and *k*′ denotes Henry’s law constant of H_2_ at the incubation temperature (i.e. 343.15 K) and at 298.15 K, respectively, Δ_*soln*_*H* is the enthalpy of solution and *R* is the ideal gas law constant. The constants *k*′ and 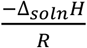 were obtained from Sander 2014 (96). Michaelis-Menten curves and parameters were estimated using the nonlinear fit (Michaelis-Menten, least squares regression) function in GraphPad Prism (version 9.3.1).

### Quantitative proteomics

Shotgun proteomics was performed to characterize the cellular adaptation and metabolic remodelling of *A. brierleyi* at different growth stages and availability of growth substrates. Quadruplicate cultures were harvested at four various conditions: (a) mid-exponential growth phase on heterotrophic medium; (b) stationary phase on heterotrophic medium; (c) transition from heterotrophic to aerobic hydrogenotrophic growth on mineral medium; (d) transition from heterotrophic to sulfur-dependent anaerobic hydrogenotrophic growth on mineral medium. Sixteen independent cultures for proteomics were first prepared by inoculating 1% v/v late exponential growing culture in 30 ml DSM medium 150 in 125 ml aerated conical flasks at incubation conditions previously described.

Cultures for conditions (a) and (b) were harvested when reaching OD_600_ values 0.04 – 0.08 pre-OD_max_ (mid-exponential growth phase) and OD_600_ values 0.04 – 0.08 post-OD_max_ (stationary growth phase), respectively. Cells maintained at autotrophic conditions (c) and (d) were prepared as follows: upon reaching a pre-ODmax OD600 of approximately 0.05, 15 ml culture was transferred into a 100kDa Amicron Ultra-15 Centrifugal Filter Unit. The tube was centrifuged at 3000 × *g* at room temperature for 5 min to separate living cells from the medium. The process was repeated once with the remaining culture. Cells on filter membrane were resuspended and washed by 3 ml of DSMZ medium 150 mineral base and the unit was centrifuged for 3 min to remove residual heterotrophic substrates. The wash step was repeated. Cells were then resuspended in 1 ml of DSMZ medium 150 mineral base and inoculated into 30 ml mineral base in a sterile 160 ml serum vial. The mineral base, per liter, was prior supplemented with 1 ml trace element solution containing (mg l^-1^) 34.4 MnSO4 • H_2_O, 50.0 H_3_BO_3_, 70.0 ZnCl_2_, 72.6 Na_2_MoO_4_ • 2 H_2_O, 20.0 CuCl_2_ • 2 H_2_O, 24.0 NiCl_2_ • 6 H_2_O, 80.0 CoCl_2_ • 6 H_2_O, 1000 FeSO_4_ • 7 H_2_O, 3.0 Na_2_SeO_3_ • 5H_2_O, and 4.0 Na_2_WO_4_ • 2H_2_O for autotrophic growth (97). For condition (c), the sealed vial was flushed with 1 bar compressed air (Industrial grade; BOC Australia) fitted with a 0.22 μm syringe filter for 2 min to establish a stable aerobic condition. 20% H_2_ and 5% CO_2_ vial headspace was prepared through ultra-high purity H_2_ and CO_2_ amendment. For condition (d), 0.3 g of tyndallized elemental sulfur powder was added to the culture and the vial was then sealed with a butyl rubber stopper. It was then flushed with 1 bar ultra-high purity N_2_ (99.999%; BOC Australia) fitted with a 0.22 μm syringe filter for 2 min to establish an anaerobic condition and amended with a final H_2_ and CO_2_ concentration of 20% and 5%, respectively. Cultures for both autotrophic conditions were allowed to grow at 70 °C and 100 rpm agitation for 48 hrs before harvesting.

To harvest cells for proteomic analysis, cell pellets from 20 ml cultures were collected by centrifugation (4800 × *g*, 50 min, −9 °C). Cell pellets were washed and resuspended with 2 ml DSMZ medium 150 mineral base in an Eppendorf tube, and centrifuged at 20200 × g at −9 °C for 10 min. For cultures maintained at condition (d), elemental sulfur powder was initially precipitated by centrifuging at 1000 rpm at −9 °C for 1 min and the supernatant was transferred to a new Falcon tube before further centrifugation to collect cells. To further remove free proteins, cell pellet was washed and resuspended in phosphate-buffered saline (PBS; 137 mM NaCl, 2.7 mM KCl, 10 mM Na_2_HPO_4_ and 2 mM KH_2_PO_4_, pH 7.4), and centrifuged again, twice. Collected cell pellets were immediately stored at −20 °C and sent to the Proteomics & Metabolomics Facility in Monash University for analysis.

The samples were lysed in SDS lysis buffer (5% w/v sodium dodecyl sulphate, 100 mM HEPES, pH 8.1), heated at 95 °C for 10 min, and then probe-sonicated before measuring the protein concentration using a bicinchoninic acid (BCA) assay kit (Pierce). Equivalent amounts of lysed samples were denatured and alkylated by adding TCEP (tris(2-carboxyethyl) phosphine hydrochloride) and CAA (2-chloroacetamide) to a final concentration of 10 mM and 40 mM, respectively, and the mixture was incubated at 55°C for 15 min. The reduced and alkylated proteins were then immobilised in S-Trap mini columns (Profiti) and sequencing grade trypsin was added at an enzyme to protein ratio of 1:50 and incubated overnight at 37°C. Tryptic peptides were sequentially eluted from the columns using (i) 50 mM TEAB, (ii) 0.2% v/v formic acid and (iii) 50% v/v acetonitrile and 0.2% v/v formic acid. The fractions were pooled and concentrated in a vacuum concentrator prior to MS analysis.

The same amount of peptides was injected into the mass spectrometer for each sample to allow accurate quantification of protein abundances and fair comparison across samples. Using a Dionex UltiMate 3000 RSLCnano system equipped with a Dionex UltiMate 3000 RS autosampler, an Acclaim PepMap RSLC analytical column (75 μm × 50 cm, nanoViper, C18, 2 μm, 100 Å; Thermo Scientific) and an Acclaim PepMap 100 trap column (100 μm × 2 cm, nanoViper, C18, 5 μm, 100Å; Thermo Scientific), the tryptic peptides were separated by increasing concentrations of 80% acetonitrile (ACN) / 0.1% formic acid at a flow of 250 nl min^-1^ for 158 min and analysed with a QExactive HF mass spectrometer (ThermoFisher Scientific). The instrument was operated in data-dependent acquisition mode to automatically switch between full scan MS and MS/MS acquisition. Each survey full scan (375 - 1575 m/z) was acquired with a resolution of 120,000 (at 200 m/z), an AGC (automatic gain control) target of 3 × 10^6^, and a maximum injection time of 54 ms. Dynamic exclusion was set to 15 seconds. The 12 most intense multiply charged ions (z ⩾ 2) were sequentially isolated and fragmented in the collision cell by higher-energy collisional dissociation (HCD) with a fixed injection time of 54 ms, 30,000 resolution and an AGC target of 2 × 10^5^.

The raw data files were analyzed with the Fragpipe software suite v17.1 and its implemented MSFragger search engine v3.4 (98) to obtain protein identifications and their respective label-free quantification (LFQ) values using standard parameters. Standard peptide modification was as follows: carbamidomethylation at cysteine residues was set as a fixed modification, as well as oxidation at methionine residues, and acetylation at protein N-terminal were set as variable modifications. For all experiments, peptides and their corresponding proteins groups were both filtered to a 1% FDR with Percolator (99). These data were further analyzed with LFQ-Analyst (100) as follows. First, contaminant proteins, reverse sequences and proteins identified “only by site” were filtered out. Also removed were proteins only identified by a single peptide and proteins that have not been identified consistently. The LFQ data was converted to log2 scale, samples were grouped by conditions and missing values were imputed using the ‘Missing not At Random’ (MNAR) method, which uses random draws from a left-shifted Gaussian distribution of 1.8 StDev (standard deviation) apart with a width of 0.3. Protein-wise linear models combined with empirical Bayes statistics were used for the differential expression analyses. The R package ‘limma’ was used to generate a list of differentially expressed proteins for each pair-wise comparison. A cutoff of the ‘adjusted p-value’ of 0.05 (Benjamini-Hochberg method) along with a |log2 fold change| of 1 was applied to determine significantly differentially abundant proteins in each pairwise comparison.

## Footnotes

### Data Availability

All study data are included in the article and/or supporting information.

## Acknowledgements

The study was supported by an ARC Discovery Project Grant DP200103074 (to C.G.), a Monash International Tuition Scholarships (to P.M.L.), an Australian Government Research Training Scholarships (to P.M.L.), a Monash Graduate Research Completion Award (to P.M.L.), a Monash Postgraduate Publications Award (to P.M.L.), and a Monash Summer Studentship (to E.T-M.). We thank Thanavit Jirapanjawat for technical support. This study used BPA-enabled (Bioplatforms Australia) / NCRIS-enabled (National Collaborative Research Infrastructure Strategy) infrastructure located at the Monash Proteomics and Metabolomics Facility.

## Author contributions

C.G. and P.M.L. conceived, designed and supervised the study. Different authors were responsible for culture preparation and maintenance (P.M.L., E.T-M. and L.J.), growth analysis (P.M.L. and E.T-M.), genome analysis (P.M.L.), phylogenetic analysis (P.M.L., C.G. and M.M.), protein structural modelling (R.G., P.M.L. and C.G.), gas chromatography measurements (P.M.L., E.T-M. and L.J.), H_2_ uptake kinetic characterization (P.M.L.), and shotgun proteomics (P.M.L., H.L., I.H., E.T., and R.B.S.). P.M.L., C.G. and R.G. analysed data and wrote the manuscript with inputs from all authors.

