## Supplementary figures and images for "Atmospheric hydrogen oxidation extends to the domain archaea"

### Figure S2

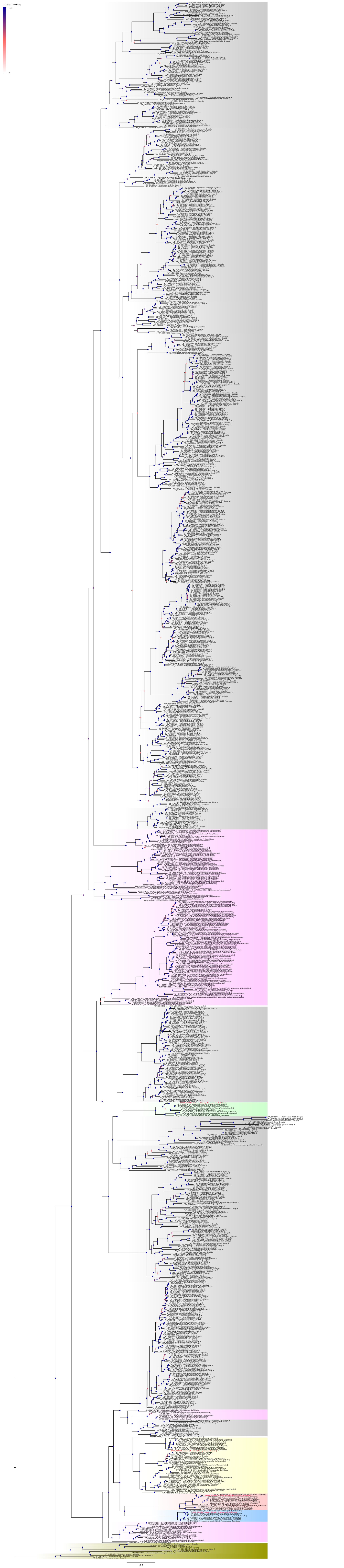
