## Supplementary information for "Atmospheric hydrogen oxidation extends to the domain archaea"

**Supplementary Figures**

**Fig. S1.** Timecourse of internal standard methane (CH_4_) concentrations in *Acidianus brierleyi* cultures and negative controls used in gas chromatography **experiments.** Headspace CH_4_ mixing ratio is presented on a logarithmic scale.

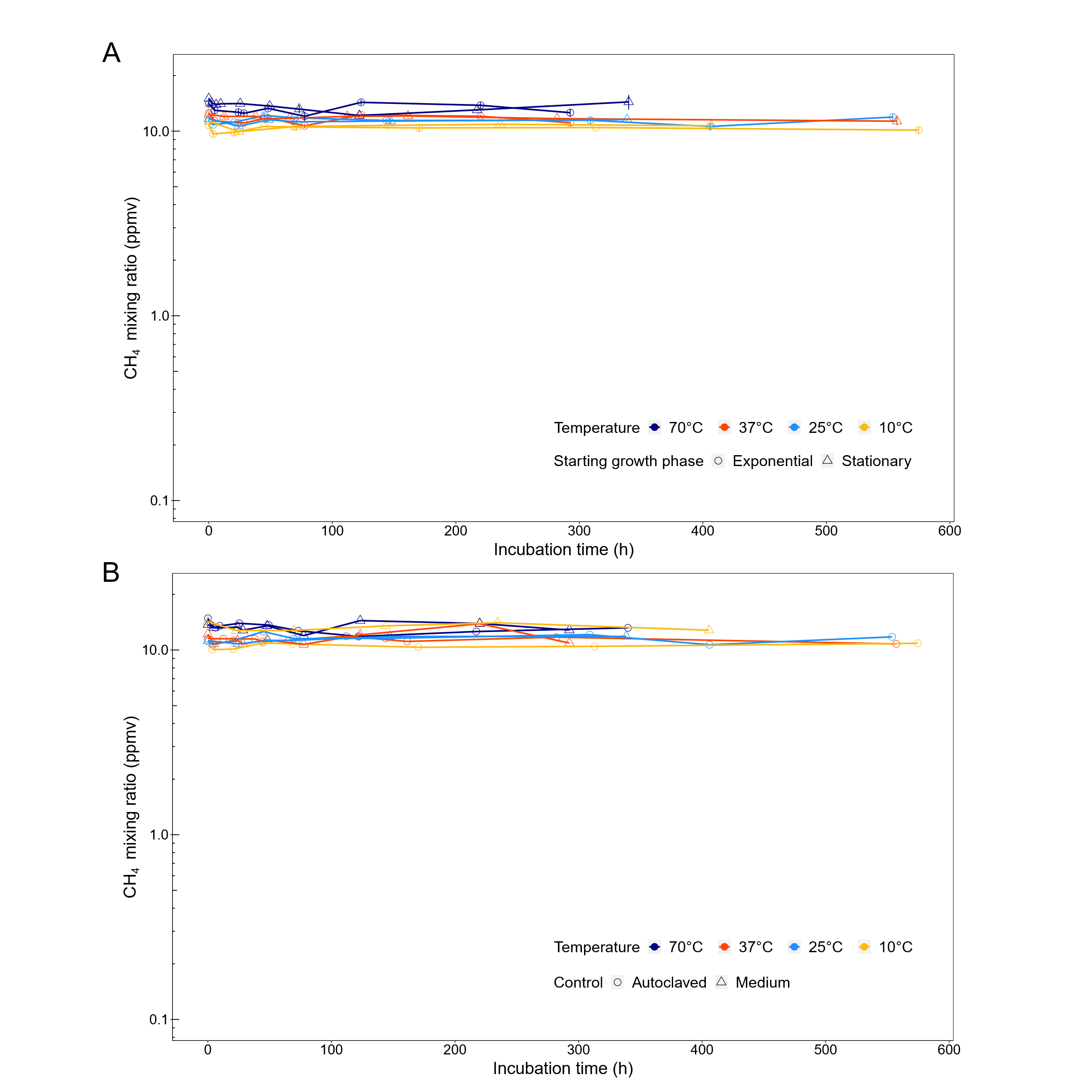

**Fig. S2.** Full maximum-likelihood phylogenetic tree of amino acid sequences of uptake group 1 and 2 [NiFe]-hydrogenases large/catalytic subunit identified in *A. brierleyi* (4 sequences; red text), genomes of all archaeal representative species in Genome Taxonomy Database (GTDB) release 202 (202 sequences), and hydrogenase reference database HydDB (1003 sequences). Group 3 and 4 [NiFe]-hydrogenases were included as outgroups and the phylogeny was rooted between group 4 [NiFe]-hydrogenases and all other groups. Subgroups/clades that were exclusively bacterial or archaeal are shaded in grey or pink, respectively. Details on alignment and tree inference can be found in **Materials and Methods** and all sequences are provided in **Table S2**. Each node was colored by ultrafast bootstrap support percentage (1000 replicates) and the scale bar indicates the average number of substitutions per site.

**Fig. S3.** AlphaFold modelling of Hsu1. (**A**) An AlphaFold model of the HsuS1 and HsuL1 subunits shown as a cartoon representation (**left panel**) and the electron transport relay formed by modelled cofactors, with predicted electron donors and acceptors indicated (**bottom panel**). (**B**) An AlphaFold model of a complex between Hsu1 and the SdhB2, SdhC2 and SdhD2 proteins, which are encoded upstream of the HsuS1 subunit. An intimate complex was not predicted between HsuS1/L1 and SdhB2/C2/D2. However, an [FeS]-cluster of both complexes is within electron transfer distance in the model suggesting SdhB2/C2/D2 may accept electrons from Hsu1 for the reduction of quinone via a transient interaction between the two complexes.

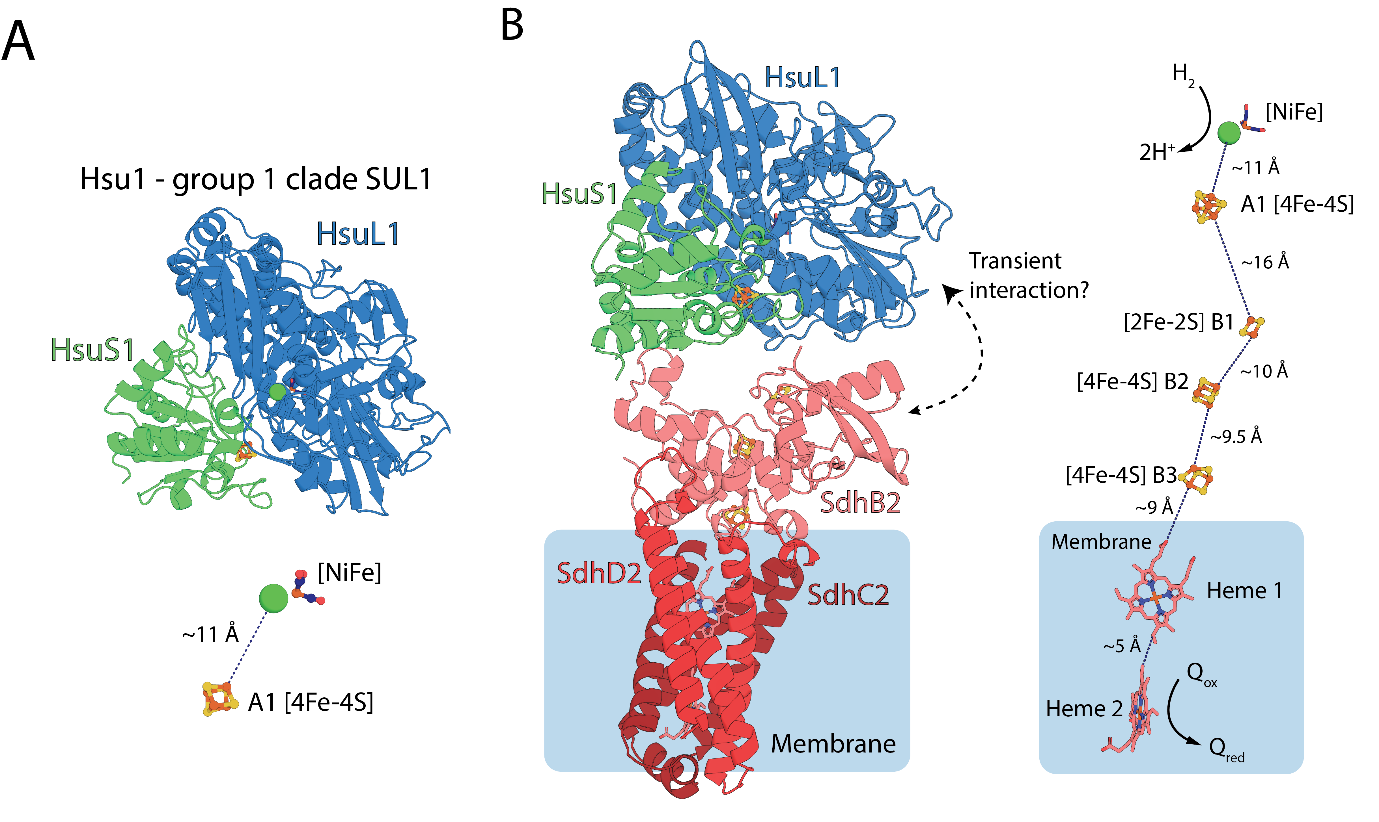

**Fig. S4.** Carbon monoxide dehydrogenase operon of *Acidianus brierleyi*.

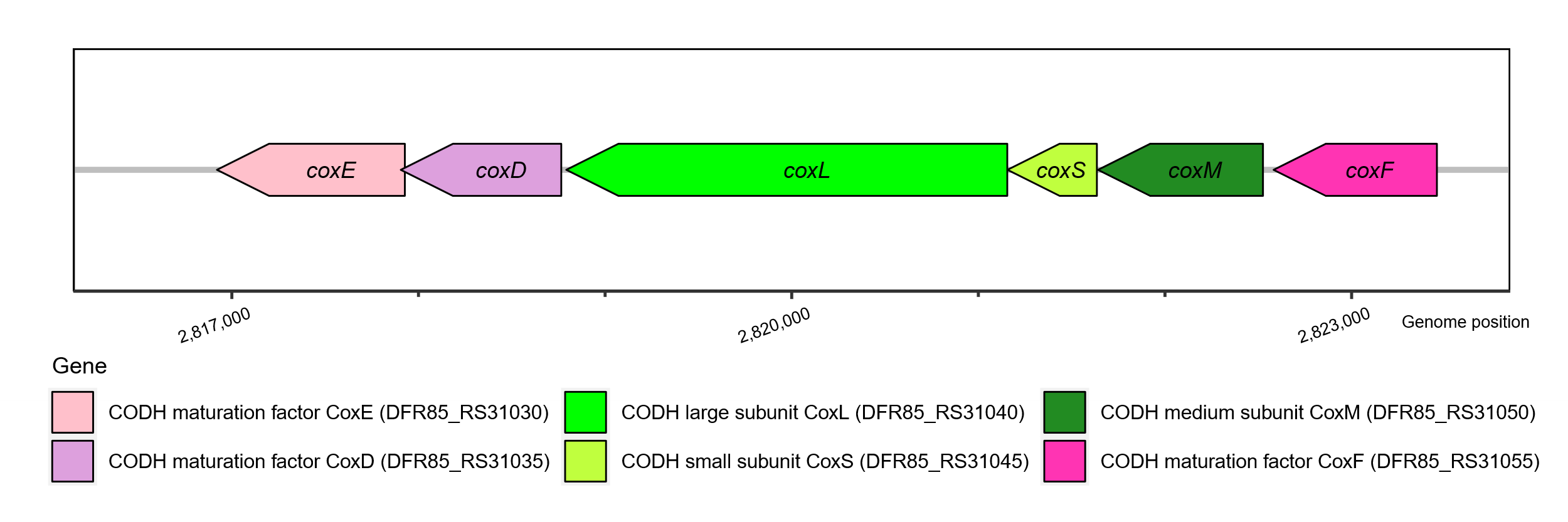

**Fig. S5.** Principal component analysis (PCA) showing distinct proteome clustering of *Acidianus brierleyi* grown at four different conditions. Mid-exponential growth on heterotrophic medium (EX); stationary phase on heterotrophic medium (ST); transition from heterotrophic to sulfur-dependent anaerobic hydrogenotrophic growth on mineral medium (AN); transition from heterotrophic to aerobic hydrogenotrophic growth on mineral medium (AE).

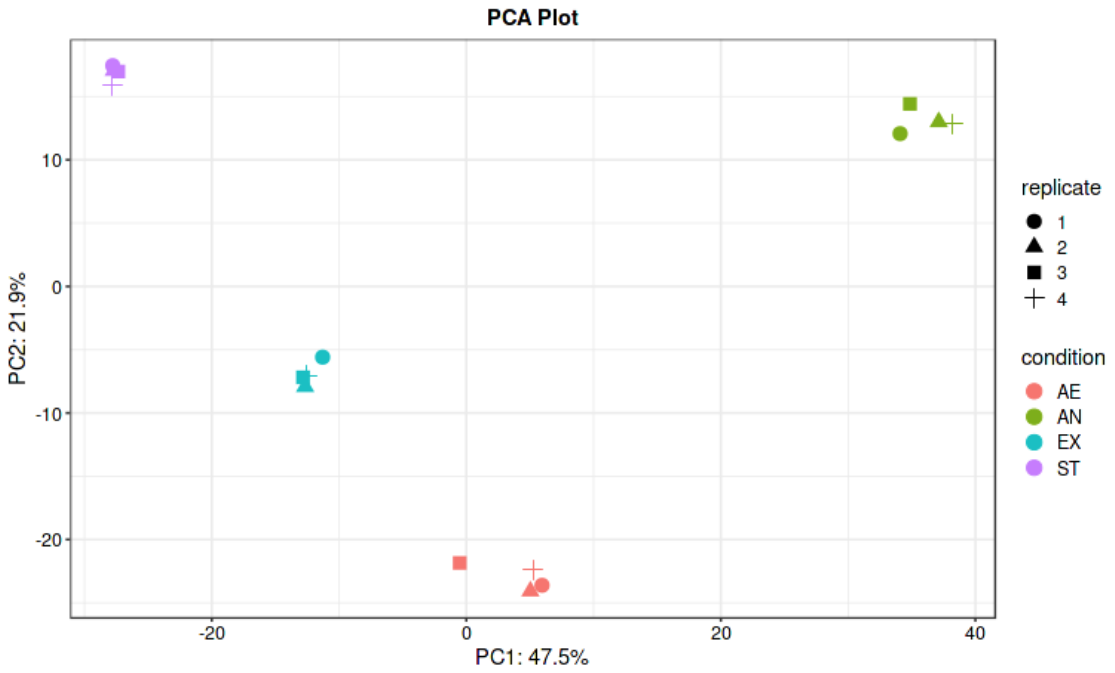

**Fig. S6.** Quantitative comparison of *Acidianus brierleyi* proteins involved in hydrogenase maturation, sulfur oxidation and iron uptake/oxidation under heterotrophic growth, stationary phase, sulfur-dependent hydrogenotrophic growth and aerobic hydrogenotrophic growth. Culture condition (four biological replicates each): EX, mid-exponential growth phase on heterotrophic medium; ST, stationary phase on heterotrophic medium; AN, anaerobic sulfur-dependent hydrogenotrophic growth; AE, aerobic hydrogenotrophic growth (**Methods and Materials**). Normalised protein abundance value represents MaxLFQ total intensity for the protein. Bubble size and color indicate protein abundance of the corresponding gene product in each biological replicate. Significant difference in fold changes of protein abundance of each condition pair is denoted by asterisks (adjusted p value ≤0.001, ***; ≤0.01, **; ≤0.05, *; > 0.05, ns).

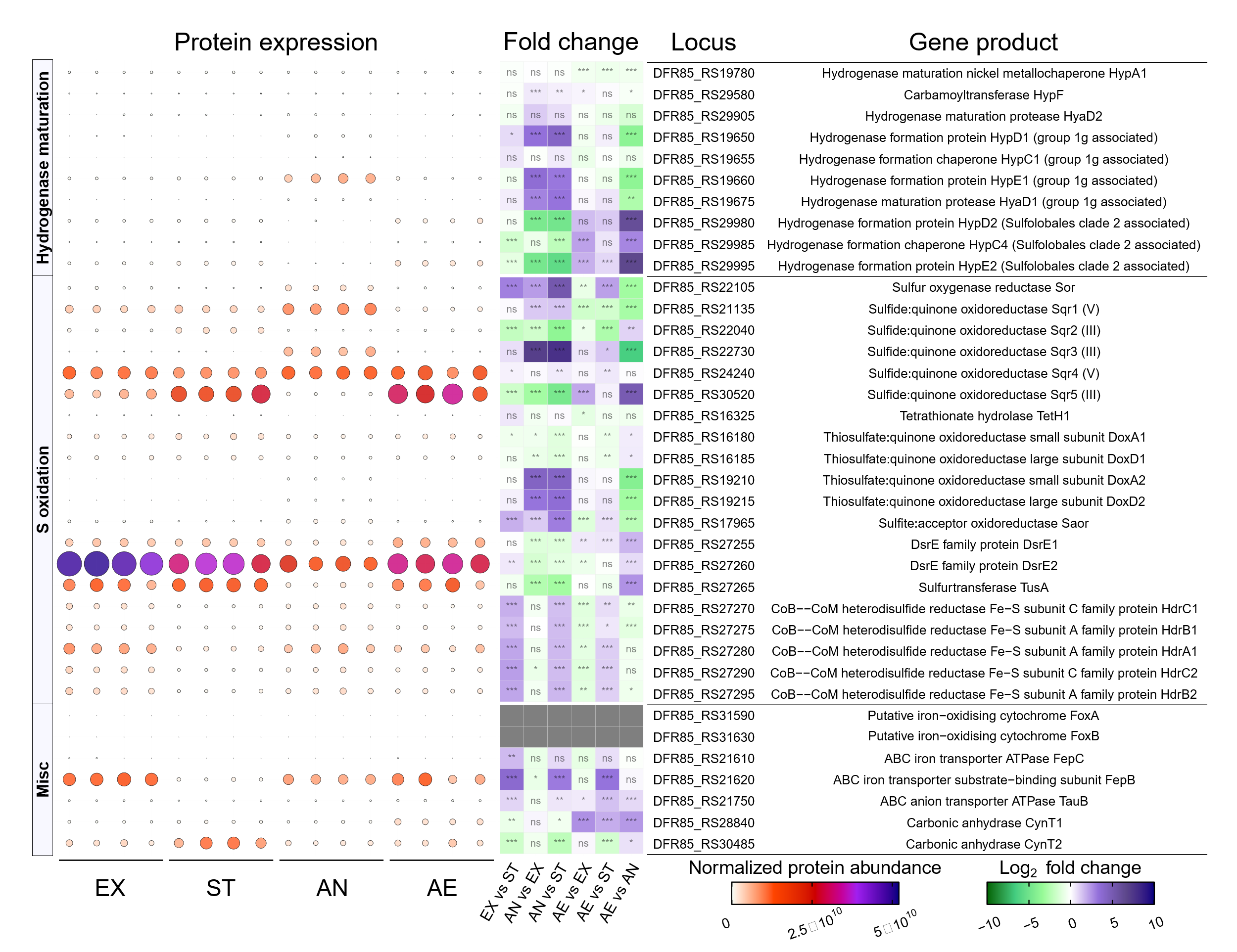

**Fig. S7.** Quantitative comparison of *Acidianus brierleyi* proteins involved in reserve degradation and amino acid/peptide/sugar/acid transport under heterotrophic growth, stationary phase, sulfur-dependent hydrogenotrophic growth and aerobic hydrogenotrophic growth. Culture condition (four biological replicates each): EX, mid-exponential growth phase on heterotrophic medium; ST, stationary phase on heterotrophic medium; AN, anaerobic sulfur-dependent hydrogenotrophic growth; AE, aerobic hydrogenotrophic growth (**Methods and Materials**). Normalised protein abundance value represents MaxLFQ total intensity for the protein. Bubble size and color indicate protein abundance of the corresponding gene product in each biological replicate. Significant difference in fold changes of protein abundance of each condition pair is denoted by asterisks (adjusted p value ≤0.001, ***; ≤0.01, **; ≤0.05, *; > 0.05, ns).

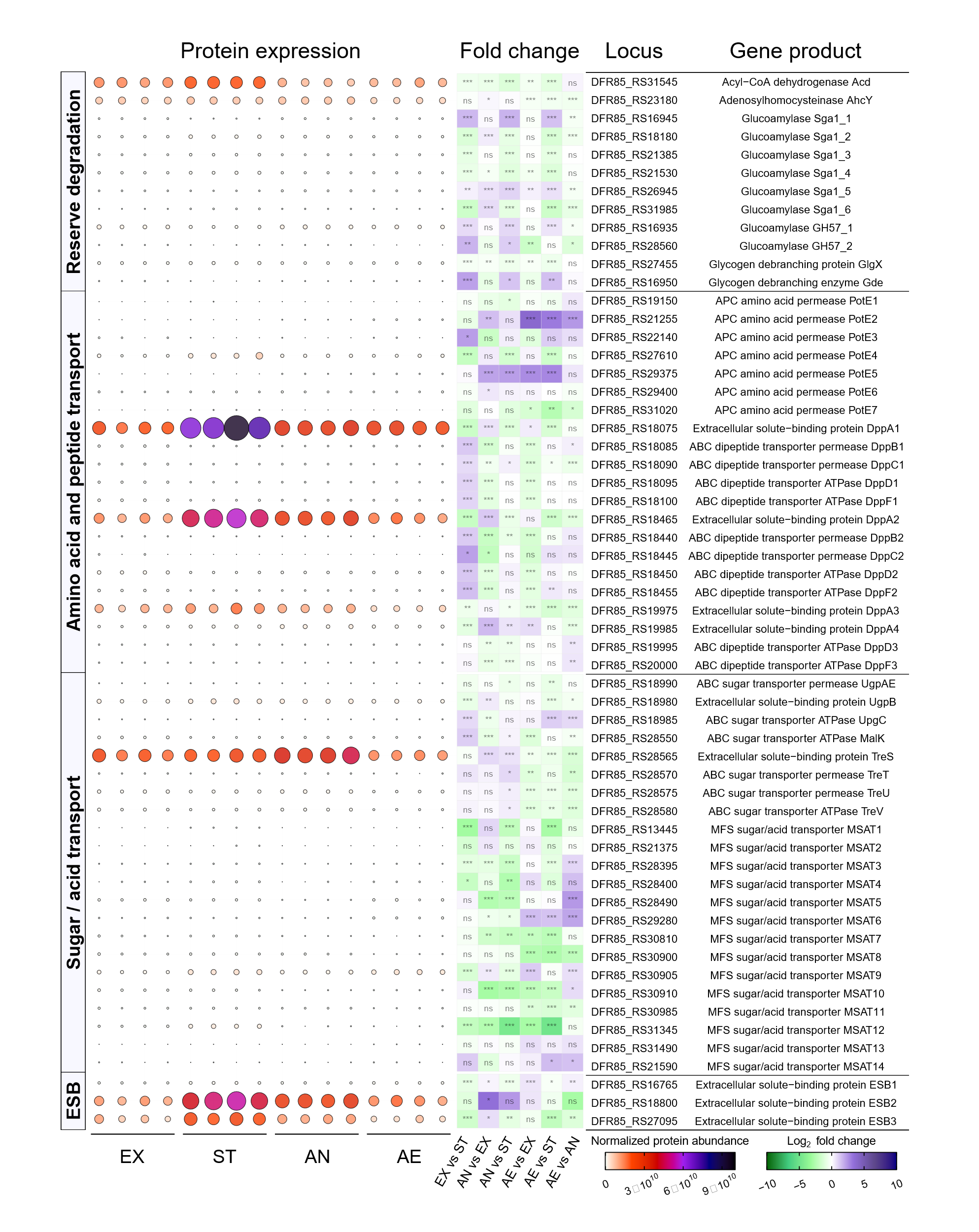

**Supplemental Note 1 – AlphaFold modelling of the *A. brierleyi* hydrogenases**

**Hca - Group 1g hydrogenase**

To generate the structural model of Hca from *A. brierleyi,* the amino acid sequences of the four putative structural subunits HcaS, HcaL, Isp1-1, and Isp2-1 were provided to AlphaFold multimer. AlphaFold convincingly assembled all four proteins into a single complex, supporting previous work indicating that HcaS, HcaL and Isp1-1 form a complex in *Acidianus ambivalens* (1). As predicted based on sequence homology, Isp1-1 is an integral membrane di-heme *b*-type cytochrome protein that forms a complex with the extracellular [NiFe]-hydrogenase catalytic subunit formed by HcaS and HcaL, which is anchored to the membrane by a transmembrane helix at the C-terminus of HcaS. Based on homology to the NarI subunit of nitrate reductase, we modelled two heme molecules in the integral membrane region of Isp1, both exhibiting bidentate coordination by conserved histidine residues (**Fig. 4B**). Interaction between Isp1 and HcaS positions one of these hemes within electron transfer distance of the distal [4Fe-4S]-cluster of HcaS, providing a route for electrons produced by H_2_ oxidation by HcaL. In hydrogenases and related enzymes, *b*-type cytochromes generally reduce or oxidize respiratory quinone, providing a link between the enzyme and the cellular electron transport chain (2–4) (**Fig. 4B**). As our proteomic data shows Hca primarily functions as a H_2_ oxidizing enzyme during anaerobic sulfur-reducing metabolism, it is likely that Isp1 reduces quinone. Intriguingly, the previously uncharacterized Isp2-1 subunit forms a complex with Isp1-1 on the cytoplasmic side of the membrane. Isp2-1 contains three iron-sulfur clusters, two of which have a cubane [4Fe-4S] configuration, and one of which resembles the non-cubane [4Fe-4S] from the HdrBC subunits of CoB-CoM heterodisulfide reductase, which Isp2-1 is structurally homologous to (5). The role of Isp2 in Hca function requires experimental verification. However, this model indicates that it provides a conduit for electrons derived from H_2_ oxidation to the cytoplasm, where they could be used to reduce NAD^+^ or ferredoxin to drive carbon fixation during autotrophic growth. The dual path for electrons within Hca creates the possibility that electron bifurcation may occur, with electrons from H_2_ split between relatively high potential quinone and a low potential electron acceptor like ferredoxin or NAD^+^.

**Hsu2 - SUL2 hydrogenase**

While Hsu2 is from a distinct phylogenetic lineage to Hca (**Fig. 3**), based on its associated genes, its subunit composition appears to be analogous to Hca, consisting of HsuS2, HsuL2, Isp1-2, and Isp2-2 subunits (**Fig. 4A**). Consistent with this, the model of Hsu2 produced by AlphaFold is similar to Hca with an extracellular membrane-anchored [NiFe]-hydrogenase module composed of HsuS2 and HsuL2 interacting with the integral membrane Isp1-2, with heme and FeS cluster distances compatible with electron transfer. As with Hca, the Isp2-2 forms a complex with Isp1-2 on the cytoplasmic side of the membrane. Isp2-2 only coordinates two [4Fe-4S] clusters, lacking the predicted cubane cluster present in Isp2-1 (**Fig. 4C**). Despite this, like Isp2-1 in Hca, Isp2-2 provides a conduit for electrons from H_2_ oxidation by Hsu to the cytoplasm, possibly for the direct reduction of substrates utilized for carbon fixation. The similarity of the structural models of Hca and Hsu2 is intuitive given their distinct expression profiles and likely allows these two proteins to fulfil analogous roles, with Hca acting under anaerobic conditions with sulfur as the electron acceptor and Hsu2 acting under aerobic conditions with O_2_ as the electron acceptor.

**Hys – Group 2e hydrogenase**

Upstream of the genes encoding the Hys [NiFe]-hydrogenase catalytic subunits HysS and HysL, and encoded in the opposite direction, is a 165 amino acid protein distantly related to the HucM subunit of Huc, a group 2a hydrogenase produced by *Mycobacterium smegmatis* (**Fig. 4A**). Recent structural analysis of Huc shows that HucM forms a tetramer that acts as a scaffold, forming a complex with four dimers of the HucSL [NiFe]-hydrogenase catalytic subunit. Each catalytic subunit is composed of HucS and HucL proteins, with the Huc complex containing four HucM, eight Huc and eight HucS subunits (6). Based on this arrangement, we hypothesized that the Hys catalytic subunits form a dimer, which was confirmed by the AlphaFold model produced when the program was provided with two HysS and two HysL subunits. This structure is highly similar to the catalytic dimer of HucSL (RMSD = 1.55 Å for 5613/8158 atoms) and contains conserved amino acids responsible for coordinating menaquinone at the Huc electron acceptor site (**Fig. 4D**). Interestingly, when AlphaFold was provided with four HysM subunits, it produced a tetrameric coiled-coil tube that is topologically and structurally similar to HucM (**Fig. 4E**).  Moreover, like HucM, the HysM tube is exclusively lined with hydrophobic residues. As well as serving as a scaffold for the Huc complex, the end of the HucM tube interacts with the cytoplasmic membrane, acting as a conduit allowing hydrophobic menaquinone to reach the electron acceptor site in HucS (6). The analogous structure and hydrophobic nature of the HysM tube suggest that it plays a similar role in Hys, which is remarkable considering the distant relationship between archaeal Hys and bacterial Huc. Unlike HucM, the AlphaFold model of HysM forms a compact tube, which lacks an obvious region that would facilitate the binding of Hys catalytic dimers. However, this may be a limitation of the AlphaFold modelling, as due to memory constraints we were not able to model all subunits concurrently. Alternatively, an additional unidentified subunit may be required to scaffold the complex. However, no candidates were identified by AlphaFold modelling of protein encoded by genes in proximity to HysS or HysL. Based on the similarity of Hys to Huc, we have generated a plausible model of the complete Hys complex (**Fig. 4E**).

**Hsu1 – SUL1 hydrogenase**

The Hsu1 gene cluster contains genes encoding the [NiFe]-hydrogenase catalytic HsuS1 and HsuL1 subunits (**Fig. 4A**). HusS1 is truncated compared to other [NiFe]-hydrogenase small subunits and, as a result, only contains a single [4Fe-4S]-cluster (**Fig. S3A**). Based on these subunits, it is unclear what the electron carrier for Hsu1 is. However, upstream of the HsuS1 gene are the three open reading frames encoding the proteins SdhB2, SdhC2, and SdhD2 (a gene encoding SdhA-like protein was not identified next to the three genes), homologous to the three subunits of succinate dehydrogenase. In succinate dehydrogenase, these subunits form a [FeS]-cluster and heme-containing integral membrane complex that relays electrons between the succinate oxidizing subunit and membrane-bound quinone (7). It is possible that these proteins serve as the electron relay from the Hsu1 catalytic subunit, meaning that Hsu1 also reduces quinone. However, while AlphaFold modelling placed the subunits within plausible electron transfer distance, a stable interface between the HsuS1/HsuL1 and SdhB2/SdhC2/ SdhD2 was not predicted (**Fig. S3B**). This makes it unclear if these proteins are functionally related.

**Supplementary Tables**

**Table S1 (xlsx). Gas chromatography measurements of H_2_, CO, and CH_4_ concentrations in tested cultures and negative controls.**

**Table S2 (xlsx). Group 1 and 2 [NiFe] hydrogenases used for phylogenetic analysis.**

**Table S3 (xlsx). Annotations and protein expressions of predicted genes of *Acidianus brierleyi*.**
